# Systematic analysis of low-affinity transcription factor binding site clusters *in vitro* and *in vivo* establishes their functional relevance

**DOI:** 10.1101/2021.12.17.473130

**Authors:** Amir Shahein, Maria López-Malo, Ivan Istomin, Evan J. Olson, Shiyu Cheng, Sebastian J. Maerkl

## Abstract

Transcription factor binding to a single binding site and its functional consequence in a promoter context are beginning to be relatively well understood. However, binding to clusters of sites has yet to be characterized in depth, and the functional relevance of binding site clusters remains uncertain. We employed a high-throughput biochemical method to characterize transcription factor binding to clusters varying across a range of affinities and configurations. We found that transcription factors can bind concurrently to overlapping sites, challenging the notion of binding exclusivity. Furthermore, compared to an individual high-affinity binding site, small clusters with binding sites an order of magnitude lower in affinity give rise to higher mean occupancies at physiologically-relevant transcription factor concentrations *in vitro*. To assess whether the observed *in vitro* occupancies translate to transcriptional activation *in vivo*, we tested low-affinity binding site clusters by inserting them into a synthetic minimal CYC1 and the native PHO5 *S. cerevisiae* promoter. In the minCYC1 promoter, clusters of low-affinity binding sites can generate transcriptional output comparable to a promoter containing three consensus binding sites. In the PHO5 promoter, replacing the native Pho4 binding sites with clusters of low-affinity binding sites recovered activation of these promoters as well. This systematic characterization demonstrates that clusters of low-affinity binding sites achieve substantial occupancies, and that this occupancy can drive expression in eukaryotic promoters.

## Introduction

DNA regulatory sequences, especially in eukaryotic organisms, frequently contain clusters of proximal binding sites (Gotea et al. (2010); Ezer et al. (2014a); Lifanov et al. (2003)). Both theory and experiment suggest that this clustering of binding sites is necessary for effective regulatory function (Wunderlich and Mirny (2009); Estrada et al. (2016); Gotea et al. (2010)). Accordingly, binding site clusters, in particular those composed of binding sites for the same transcription factor (homotypic clusters) or of low-affinity sites, have received increasing attention with several functionally important examples identified to date (Crocker et al. (2015); Kribelbauer et al. (2019); Crocker et al. (2016)). For instance, clusters of low-affinity binding sites were found to be critical for the precise temporal control of gene expression (Gaudet and Mango (2002)), spatial control and patterning in development (Crocker et al. (2015); Farley et al. (2015); Jiang and Levine (1993)), as well as robustness to mutations (Crocker et al. (2015)), features that are hallmarks of native biological systems and highly relevant for engineering synthetic systems. However, our understanding of how such outcomes can be encoded in eukaryotic regulatory sequences and transduced by transcription factors, co-factors, and other biological machinery into function is still in its infancy. In prokaryotes (Wunderlich and Mirny (2009); Whyte et al. (2013); Estrada et al. (2016); de Boer et al. (2020)) and eukaryotes (Rajkumar et al. (2013); Segal et al. (2008); Mogno et al. (2013)), biophysical models have had success in predicting gene expression based on sequence information alone, but we are missing comparable insights where DNA regulatory sequences contain complex clusters of binding sites. In order to build a better understanding of eukaryotic transcription, we argue that it is important to characterize and develop a quantitative understanding of how transcription factors bind to multiple proximal binding sites varying in affinity, multiplicity, and density (Weingarten-Gabbay and Segal (2014)).

The bulk of existing research on TF-DNA interactions has revolved around single binding sites, often focusing on consensus sequence or high-affinity binding sites. We currently lack a detailed, biochemical analysis of the binding of transcription factors to binding site clusters. For instance, while several reports identified that transcription factors act synergistically when binding to clusters of low-affinity sites (Segal et al. (2008); Castellanos et al. (2020)), whether this synergy is a biochemical property of the TF-DNA interaction or depends on additional *in vivo* complexity remains unknown. Current biochemical approaches for studying transcription factor-DNA interactions either do not provide a direct readout of the occupancy of multiple transcription factors interacting on DNA (Levo et al. (2015); Zhu et al. (2018)), or lack the throughput necessary to tackle the larger sequence space that emerges when working with multiple sites (Kim et al. (2013); Castellanos et al. (2020)). Furthermore, current methods tend to be biased towards high-affinity interactions, due to the process of isolating bound molecules which can lead to dissociation of lower-affinity, transient binding events.

To fill this knowledge gap, we applied the quantitative, high-throughput MITOMI assay (Maerkl and Quake (2007)) to study binding site clusters. The MITOMI assay performs hundreds to thousands of parallel interaction measurements on a single microfluidic device by surface immobilizing transcription factors exposed to solution phase short DNA targets. MITOMI has been applied to the detailed quantitative characterization of protein - DNA (Maerkl and Quake (2007, 2009); Rockel et al. (2013); Blackburn et al. (2015); Le et al. (2018); Fordyce et al. (2010); Aditham et al. (2021)), protein - RNA (Martin et al. (2012)), and protein - protein interactions (Gerber et al. (2009)), as well as small molecule drug discovery (Einav et al. (2008)). The MITOMI method has also been extended to enable the high-throughput analysis of molecular association and dissociation kinetics (Geertz et al. (2012); Bates and Quake (2009)).

We slightly modified the original MITOMI assay by inverting the assay geometry using surface immobilized 90 bp-long target DNA strands and solution-phase transcription factors in order to optimize the quantitative characterization of binding site clusters. Reconfiguring MITOMI in this way allows for regulatory sequences ranging from single binding sites to large complex clusters to be encoded on DNA, and for the equilibrium occupancy of multiple transcription factor molecules to be observed. Given the throughput of our assay, we designed a library of DNA sequences that enabled direct assessment of how transcription factor binding is affected by changes to cluster configuration, including site affinities, multiplicities, and densities. We assessed transcription factors from two of the largest transcription factor families: zinc fingers and basic helix loop helix (bHLH).

We found that when DNA targets contain binding sites which overlap and share common basepairs, binding is only partially reduced compared to non-overlapping sites, suggesting that transcription factor molecules can withstand significant steric clash and simultaneously occupy two sites sharing basepairs. This effect, occupancy despite clash, decreased as the cluster density increased to require greater levels of steric clash in states where transcription factors bind neighboring sites, or as transcription factor occupancy on DNA became less energetically favorable due to lower site affinities. Binding to low-density clusters where sites do not share neighboring basepairs appeared largely independent, lacking prominent levels of binding synergy.

We also were able to show that, compared to an individual high-affinity binding site, clusters containing as few as three weak binding sites each an order of magnitude lower in affinity than the consensus sequence will reach greater occupancy levels *in vitro* at transcription factor concentrations that likely occur *in vivo*. Although clusters of weak binding sites achieved high-occupancies *in vitro*, it remained unknown whether occupancy from low-affinity, transient binding events would translate to an *in vivo* setting and give rise to functional regulatory elements. For instance, many past reports *in vivo* have considered dwell times to be an important factor determining functional gene regulation (Popp et al. (2021); Clauß et al. (2017); Callegari et al. (2019); Lickwar et al. (2012); Loffreda et al. (2017)). To determine the functional relevance of weak binding site clusters *in vivo* we generated and characterized a synthetic library of minCYC1 promoters containing multiple Zif268 binding sites, and were able to show that the gene expression level driven by clusters of low-affinity sites can match those achieved by high-affinity, consensus binding sites. Finally, to determine whether clusters of weak binding sites could function in a native gene regulatory network we tested weak binding site clusters in the inorganic phosphate regulatory network. We replaced native binding regions containing high-affinity Pho4 and Pho2 binding sites with clusters of low-affinity Pho4 sites in the PHO5 promoter and found that in the context of this native regulatory system controlled by physiological levels of Pho4 transcription factor, gene expression can be recovered with binding site clusters consisting of individual binding sites an order of magnitude lower in affinity than the consensus site.

## Results

### iMITOMI Development and Characterization

We developed an inverted MITOMI assay (iMITOMI), by adapting the original MITOMI platform (Maerkl and Quake (2007)) to generate quantitative measurements of transcription factors binding at equilibrium to longer DNA targets containing binding site clusters of up to 6 distinct binding sites (Figure 1A-H) (Istomin (2020)). We also tested and characterized these binding site clusters *in vivo* in the eukaryotic model organism *S. cerevisiae* (Figure 1I-J).

**Figure 1:**
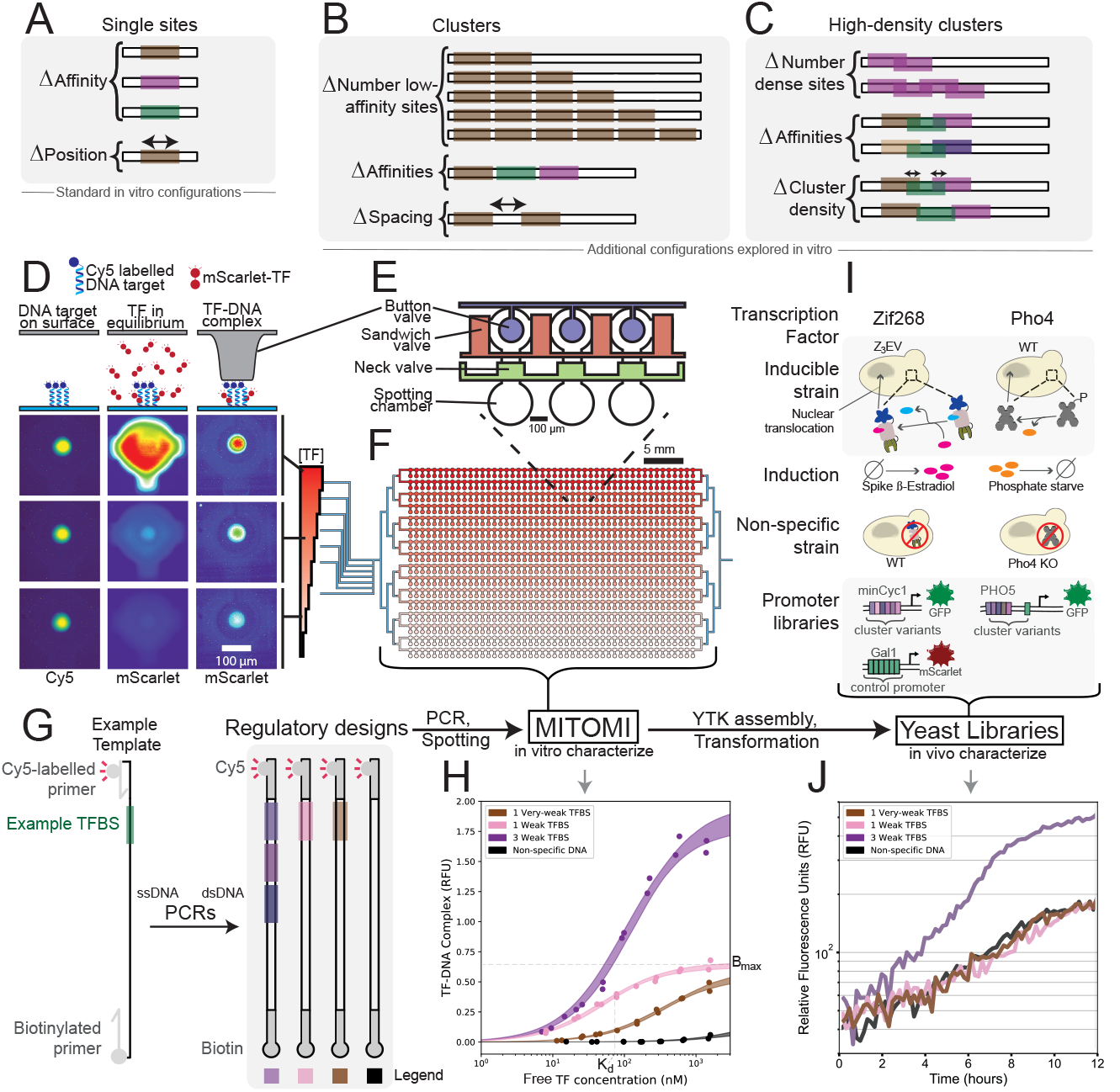
Description of the investigation and experimental process. **(A-C)** DNA libraries and parameters explored *in vitro* using iMITOMI, including clusters of multiple binding sites **(B)**, and high-density clusters of overlapping binding sites **(C). (D-F)** Schematic of the high-throughput, *in vitro* iMITOMI method. The iMITOMI chip consists of two-layers, where microvalves in the control layer **(E)** control the flow of fluid in a flow-layer containing 768 programmable reaction chambers **(F). (D)** Image rows represent different TF concentrations, columns represent stages of the assay. DNA is immobilized on the chip’s surface, in each assay chamber, under the button valves (left column). Fluorescently-tagged transcription factor molecules are flowed at different concentrations (red color gradient) into different rows of the chip. The free (middle column) and bound (right column) transcription factor signal is quantified at equilibrium. **(G)** To obtain the dsDNA targets, 90 bp-long single-stranded templates are amplified with biotinylated and Cy5-tagged primers. **(H)** Example 2-parameter saturation binding curves for DNA targets containing from 0 to 3 binding sites spanning a wide affinity-range. Shaded regions represent the 5 to 95% confidence intervals identified by exploring model parameter space with a Markov Chain Monte Carlo sampler. **(I)** *In vitro*-characterized clusters are assembled into expression cassettes, and chromosomally integrated into *S. cerevisiae* for characterization **(J)**. Zif268 regulatory sequence designs were characterized in the minCYC1 promoter using an inducible Z_3_EV system, where a wildtype (WT) strain enabled quantification of the non-specific response **(I, left)**. Pho4 regulatory sequence designs were characterized in the PHO5 promoter, in a WT strain induced by phosphate starvation, where a Pho4 knockout strain enabled quantification of the non-specific response **(I, right)**.

In our assay, 90 bp-long double stranded DNA (dsDNA) is immobilized on the chip’s surface, whereas protein is maintained free in solution and titrated across different regions of the chip (Figure 1D-H, Supplementary Figure 1). This is in contrast to the original MITOMI assay in which transcription factor was immobilized to the surface and 30 bp-long target DNA was added in solution. Inverting the assay chemistry was important to study binding site clusters for several reasons. Using surface-immobilized DNA allows for multiple transcription factor molecules to bind and the resulting increase in fluorescence to be measured, whereas binding of multiple immobilized transcription factors to individual free molecules of DNA would present spatial constraints, and could lead to skewed affinity measurements due to avidity.

The iMITOMI chip contains 768 unit cells, each consisting of a DNA spotting chamber, into which DNA targets are introduced by spotting during fabrication, and a MITOMI detection area, wherein binding occurs and detection is conducted (Figure 1D-F). The spotting chamber and detection area can be separated from each other by actuating a neck valve, and a sandwich valve can be actuated to isolate the individual reaction chambers. A circular MITOMI button valve lies above each detection area, and can be used to mechanically trap molecules bound to the surface in the detection area when this valve is pressurized.

The DNA in each unit cell is individually programmable with a DNA spotter, and fluid flow to each row of unit cells in the chip can be controlled through actuation of a combination of eight multiplexing valve control lines. We first generate a specific surface chemistry, and immobilize Cy5-tagged dsDNA from each spotting chamber under the button valve in the corresponding detection area before imaging the chip to quantify the surface-bound DNA (Figure 1D). We then flow recombinant mScarlet-tagged transcription factor into the chip (Supplementary Figure 2), using a different concentration in each of the eight pairs of rows. After the DNA and protein are allowed to bind and reach equilibrium, we image the chip to quantify the concentration of free protein at equilibrium in each chamber. Then, by actuating the button valve to isolate DNA-bound protein, and after washing away free protein, we image the chip once more to quantify the amount of protein that was in complex with DNA at equilibrium. We can thus directly measure the equilibrium concentrations of free as well as bound protein. We then use these measurements to generate binding curves (Figure 1H) by relating the bound protein signal (normalized by the DNA signal) to the free protein signal at equilibrium. For each DNA target, a Markov-Chain Monte Carlo sampler is used to fit a 2-parameter saturation binding curve model.

To validate our method, we conducted several experiments to ascertain that the affinities and specificities measured with iMITOMI correspond to previous MITOMI measurements, which were validated against data obtained with other methods generally considered a gold standard (Rastogi et al. (2018); Maerkl and Quake (2007); Geertz et al. (2012)). Affinity and specificity measurements taken for single binding site targets were found to be consistent with previous results obtained using standard MITOMI across a wide affinity range, both for Zif268 (Swank et al. (2019); Blackburn et al. (2015)) and Pho4 (Maerkl and Quake (2007)) (Supplementary Figure 1). To test whether using surface-bound DNA might influence transcription factor binding, we placed single binding sites for Zif268 and Pho4 in different positions of the DNA target. No difference in either *K*_D_ or *B*_max_ was observed as the binding site is moved closer towards the chip surface (Supplementary Figure 3), indicating that the distance from the chip surface did not introduce systematic differences in the ability of a transcription factor to bind to the target DNA strand.

### *In vitro* characterization of binding site clusters

We applied our method to characterize the binding of Pho4 from the bHLH family and Zif268 from the zinc finger family to DNA targets composed of multiple binding sites, ranging from from 1 to 6 sites for Zif268 (Figure 2A-E) and from 1 to 5 sites for Pho4 (Figure 2F-J). We characterize compositions of weak sites (W) and very-weak sites (V), in the range of one, or two orders of magnitude lower affinity than the consensus site, respectively.

**Figure 2:**
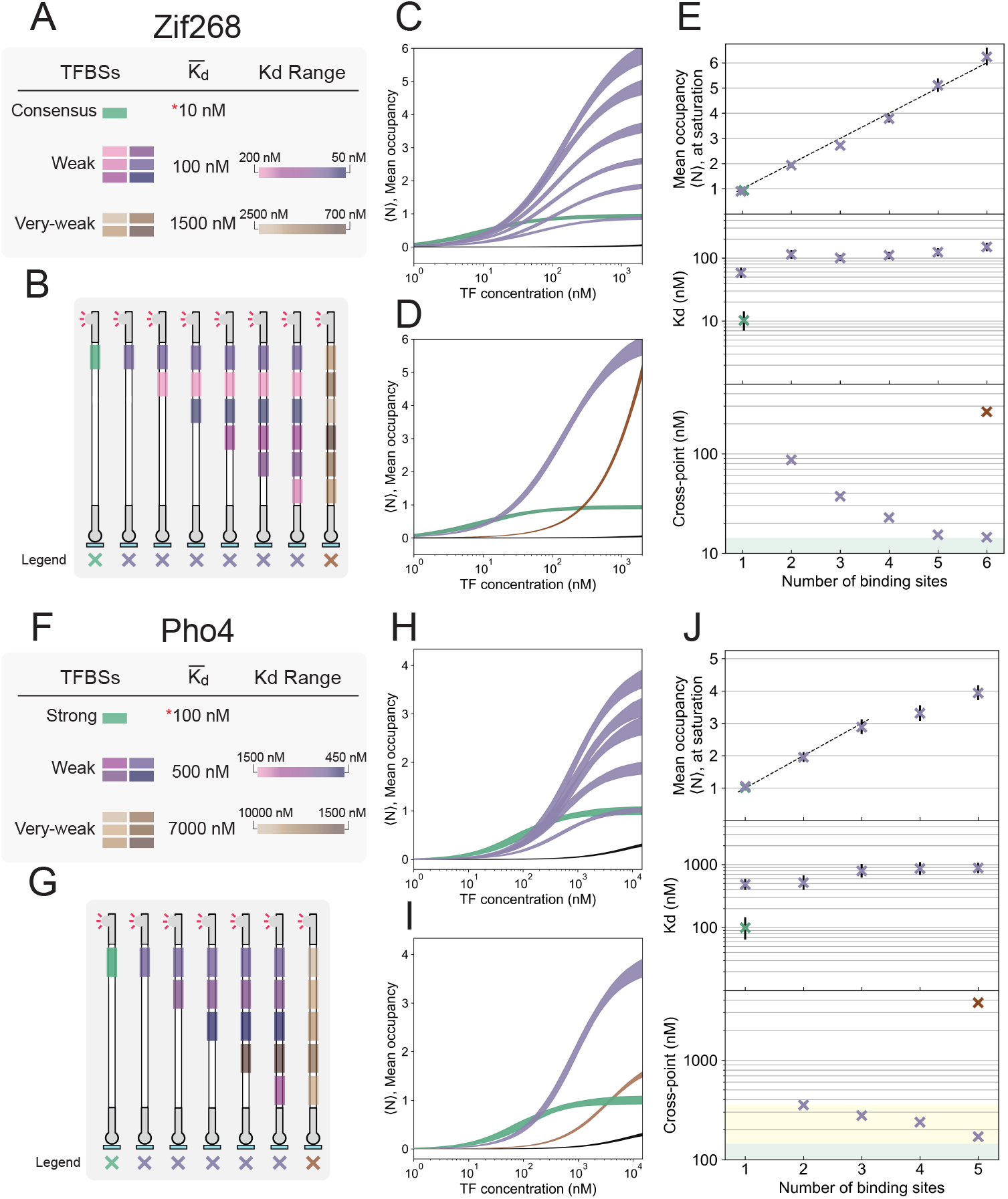
**(A, F)** Legend for the Zif268 and Pho4 clusters analyzed. Binding sites were grouped into different affinity classes. Consensus or strong sites, weak sites approximately an order of magnitude lower affinity, and very-weak sites approximately two orders of magnitude lower affinity than the consensus site. **(B, G)** Schematic of the Zif268 and Pho4 DNA targets characterized. **(C, H)** Amount of TF in complex with DNA as a function of free TF concentration at equilibrium (saturation binding curves) for DNA targets containing from 1-6 (Zif268, **(C)**) or 1-5 (Pho4, **(H)**) low-affinity binding sites (purple) compared against 1 strong binding site (green). Shaded regions represent 5 to 95% confidence intervals. Further data and analysis available in Supplementary Figure 4. **(D, I)** Saturation binding curves comparing the largest cluster of very weak sites (brown) against the largest cluster of weak sites and a single strong binding site. **(E, J, top)** Mean occupancies at saturating transcription factor concentrations, ⟨*N*⟩_max_ (*R*^2^ of 0.99 for Zif268, *R*^2^ of 0.95 for Pho4). **(E, J, middle)** Measured affinity values (*K*_D_). **(E, J, bottom)** Cross points: the transcription factor concentration at which the mean occupancy of a DNA target composed of multiple medium or weak binding sites reaches the same mean occupancy as the consensus or strong binding site target. The yellow shaded region represents a range of expected *in vivo* transcription factor concentrations, and the green shaded region represents the characterized *K*_D_ for the strong binding site. Error bars represent 5 to 95% confidence intervals.

For each transcription factor, the bound transcription factor signal (normalized by DNA fluorescence) at saturating transcription factor concentrations (which we refer to as the saturation fluorescence, *B*_max_) was similar for single strong, medium, and weak binding sites (Supplemental Figure 3). As the number of weak binding sites in a DNA target is increased, we observe a linear increase in the saturation fluorescence (*R*^2^ of 0.99 for Zif268 and 0.95 for Pho4), with a step size corresponding to the saturation fluorescence for single binding site targets (Supplementary Figure 4). This suggests that the binding sites are saturating with transcription factor, and that the step increase in *B*_max_ (for Zif268, the slope of the line in Supplemental Figure 4E) corresponds to the fluorescence signal resulting from an occupancy of 1 TF bound per DNA molecule (units of RFU per TF molecule bound on average). Therefore normalizing by this value converts our bound transcription factor signal (RFU) into a measure of mean occupancy, ⟨*N* ⟩, which is simply the average number of transcription factors bound per molecule of DNA (Figure 2C-E, H-J). In the Pho4 DNA library, the fourth binding site we introduced was a very-weak site (Supplemental Figure 3), and thus not fully saturated in the assay. Accordingly, in this case we fit our linear regression to the targets ranging from 1 to 3 binding sites to obtain the step increase in saturation fluorescence for normalization (Figure 2J).

Our results show that binding to clusters of sites an order of magnitude weaker than the consensus binding sequence can be saturated, and that all binding sites in even closely-spaced clusters can be concurrently occupied by transcription factors, resulting in a maximum occupancy ⟨*N*_max_⟩ that consistently increases by 1 with each additional site in a DNA target (Figure 2E, J, upper panels). In contrast, the measured affinity (*K*_*D*_) is determined by the individual binding site affinities present in the target cluster (Figure 2E, J, middle panels).

Having shown that clusters of weak and very-weak binding sites can be fully occupied at high transcription factor concentrations of 1 *µ*M, we next asked whether weak and very-weak clusters also lead to sufficient occupancy at lower transcription factor concentrations. The *in vivo* nuclear concentration of Pho4 in *S. cerevisiae* has previously been estimated to be approximately 355 nM based on single-cell fluorescence experiments (Rajkumar et al. (2013)). Estimates for zinc-fingers in yeast range from 538 to 3334 copies per cell, which translates to a nuclear concentration of roughly 213 nM to 1322 nM (Biggin (2011); Alberts et al. (2002)). As single consensus sequence targets are known to be physiologically functional in a cellular milieu, we determined the concentrations at which low-affinity binding site clusters exhibit similar occupancy levels as the corresponding single consensus sequence (Figure 2E, J, lower). In other words, we plotted the concentrations at which the low-affinity cluster saturation binding curves cross the single consensus target saturation binding curve. It can be readily seen that similar occupancies can be reached by all low-affinity clusters, with cross-over occurring at transcription factor concentrations of as low as 14 nM for a cluster of 6 low-affinity Zif268 binding sites, and 170 nM for a cluster of 5 low-affinity Pho4 binding sites. These values are well within the range of physiological concentration estimates and indicate that low-affinity binding site clusters may achieve comparable occupancy levels as a single high-affinity consensus site *in vivo*.

We exploited our platform to explore the impact of binding site proximity on binding affinity (Figure 3). First, we set out to determine whether proximal binding sites exhibit synergistic interaction as has been reported *in vivo* (Segal et al. (2008)). We constructed and characterized a library of targets with two low-affinity binding sites, where the position of one site was held constant, while the position of the second site was moved in intervals to adjust the spacing between sites (Δ). For Zif268, binding remained largely unchanged across the library, barring one unexplained outlier (Figure 3A, 5bp spacing). In the case of Pho4, we observed a decrease in ⟨*N* ⟩_max_ for gap distances centered around ∼ 3 bp, possibly indicating steric competition in this orientation of transcription factor molecules (Figure 3B). The *K*_D_ values remained largely constant as a function of distance, and corresponded to the individual binding site affinities. We therefore did not observe a large degree of cooperativity between adjacent binding sites (Supplementary Figure 4G), implying that the reported synergism may be a product of higher-order complexity *in vivo*. Together, our results suggest that the equilibrium binding we observe to different binding sites in targets composed of multiple sites is largely independent, possibly with mild steric effects at close distances.

**Figure 3:**
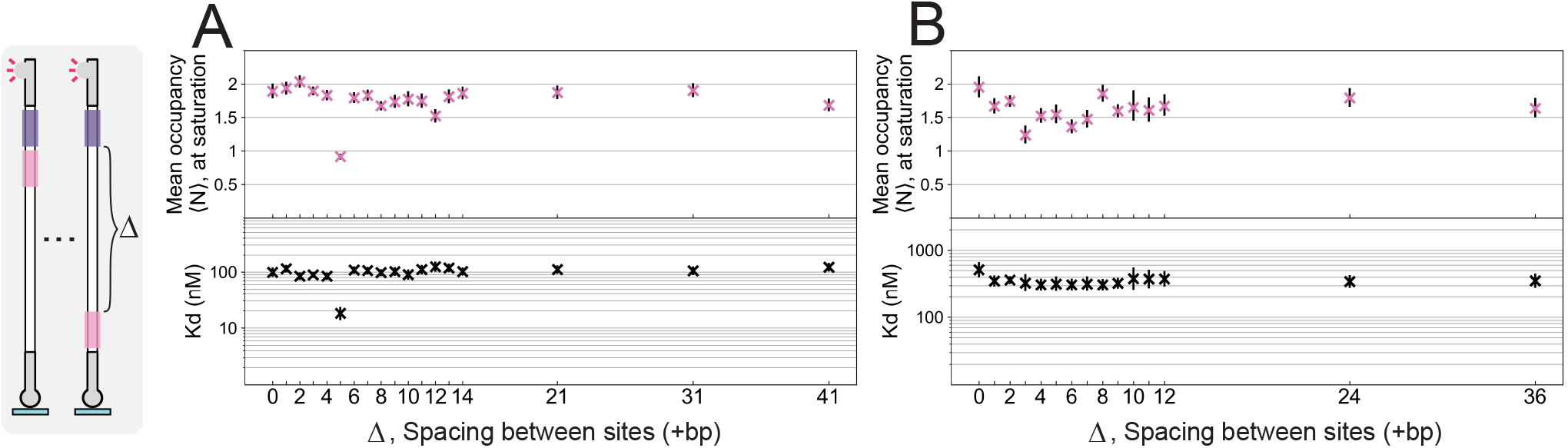
**(A, B)** Affinities (*K*_*d*_), and mean occupancies at saturation (⟨*N* ⟩_max_), for a library of DNA targets for Zif268 (A) and Pho4 (B), with different distances between two weak binding sites.

Overlapping binding sites occur frequently in regulatory DNA of prokaryotes and eukaryotes (Hermsen et al. (2006); Whyte et al. (2013); Panne (2008); Ezer et al. (2014b); de Boer et al. (2020); Hoch et al. (1992)), and are thought to influence how binding occurs among transcription factors (Segal et al. (2008); Panne (2008); Guturu et al. (2013); Fry and Farnham (1999); Stanojevic et al. (1991)), RNA polymerase, other regulatory elements (Ptashne (2011); Panne (2008); Dhiman and Schleif (2000); Humphrey et al. (1996); Lee et al. (1981)), and nucleosomes (Zhu et al. (2018)). In eukaryotes, the relatively low specificity of transcription factors gives rise to regulatory sequences that contain high-density clusters of binding sites tending to overlap with one another (Whyte et al. (2013); Panne (2008); de Boer et al. (2020)). As a result of high binding site densities, De Boer et. al. found that shifting a site by one basepair across regions of a yeast promoter impacts expression primarily through the disruption or creation of alternate transcription factor binding sites (de Boer et al. (2020)). While competition between transcription factors binding to nearby but distinct binding sites is typically thought of as a graded function of distance, as a rule of thumb overlapping binding sites are often assumed to elicit exclusive transcription factor binding due to steric clash (Fry and Farnham (1999); Lee et al. (2018)), and consequently binding models tend to ignore states where two transcription factors are bound simultaneously to sites sharing common basepairs (Segal et al. (2008)).

To our knowledge, binding of transcription factors to high-density clusters has not been characterized in detail. Thus we designed DNA targets containing from 1 to 4 binding sites of varying affinity for Zif268, where one or three basepairs are shared between neighbouring binding sites (Figure 4A). An alignment of the protein-DNA complexes to the DNA sequences together with residue-clash prediction suggests that the protein molecules will exhibit significant steric clash as binding site motifs start to overlap and share common basepairs (Figure 4B, C). Yet surprisingly, we discovered that the mean occupancy of transcription factor molecules binding to two binding sites that share basepairs can exceed a value of 1 at the upper range of physiologically relevant transcription factor concentrations (Figure 4D). This demonstrates that occupancy despite clash can occur, where two transcription factor molecules bind at once and exhibit steric interference on binding sites sharing common basepairs, and that this might occur in high-density clusters in the genome.

**Figure 4:**
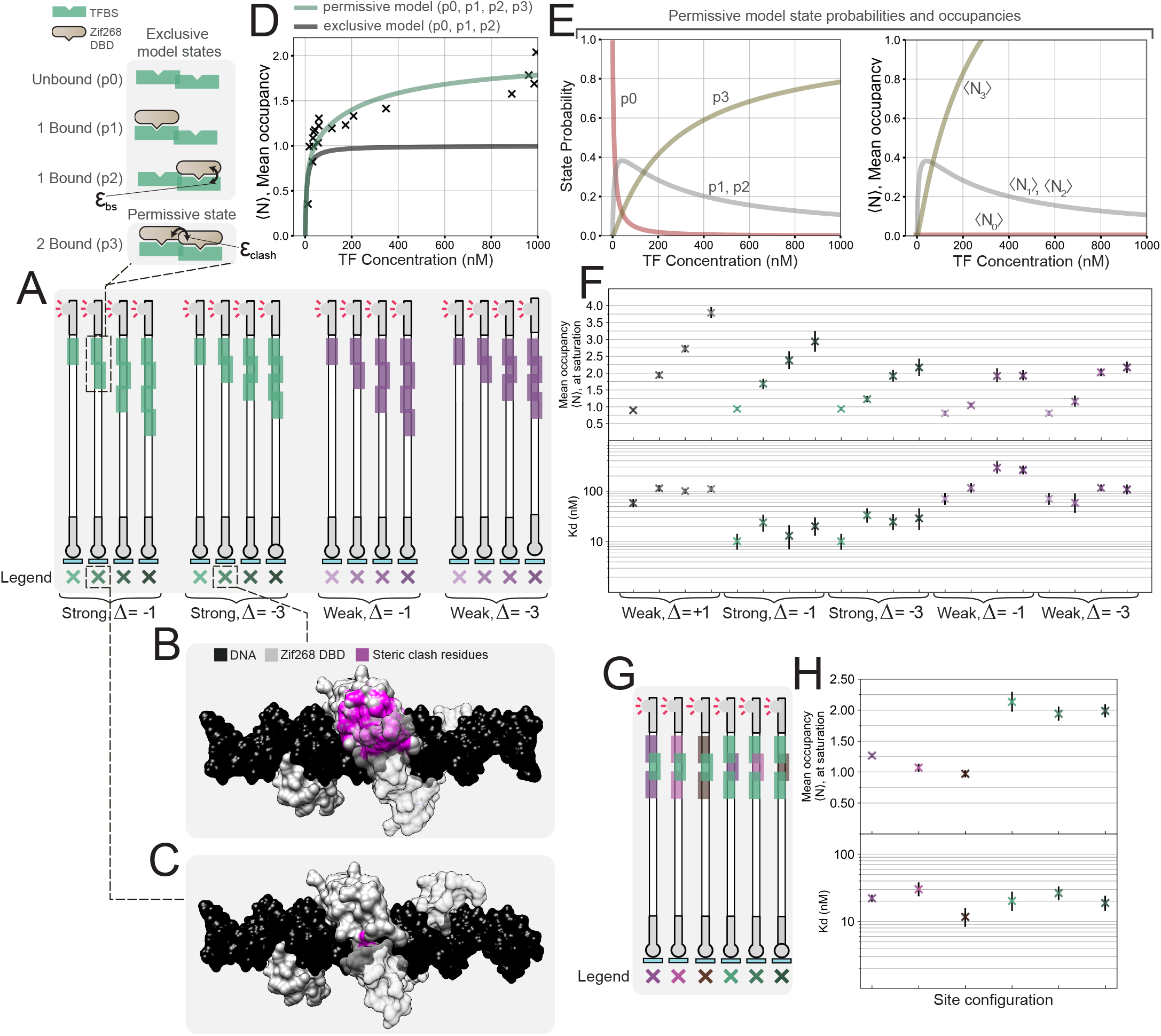
**(A)** Library of DNA targets analyzed containing Zif268 high density binding site clusters, where basepairs are shared between neighboring binding sites. Clusters contain up to four strong or weak binding sites, and neighboring sites share 1 or 3 common basepairs. **(B, C)** Aligned crystal structures, highlighting Zif268 residues expected to exhibit steric clash in magenta. 3 basepairs (33% of the Zif268 motif) shared in **(B)**, and 1 basepair (11% of the motif) shared in **(C)**). **(D-E)** Statistical mechanical analysis of binding to consensus binding sites sharing a common basepair. **(D, left)** Legend of the states corresponding to two different binding models, an exclusive binding model (top three states), and a permissive binding model (all four states). **(D, right)** Binding data and model predictions, illustrating that two transcription factors can bind at once to binding sites sharing basepairs. **(E)** Predicted probability **(left)** and occupancy **(right)** for each permissive model state as a function of transcription factor concentration. **(F)** Experimental data for the full library of high-density clusters (data fit with a 2-parameter SBC similar to above). Occupancies at saturation, ⟨*N* ⟩_max_, and affinities, *K*_*D*_. **(G)** DNA targets containing mixed-affinity high-density clusters where 3 basepairs are shared between neighboring binding sites. **(H)** Corresponding experimental data for mixed-affinity clusters.

Since a simple one-site saturation binding model fails to describe this phenomenon, we extended our description of binding to a statistical mechanical model. First we attempted to model binding to high-density clusters using an exclusive binding model (full steric occlusion) where two transcription factors cannot bind concurrently to overlapping binding sites, resulting in just three states of the system for a 2-site cluster (Figure 4D, Supplementary Table 5). We parameterized the model using binding energies (*ε*_*bs*_) derived from an independent characterization of the individual binding sites. As expected, for a two-site cluster the exclusive model fails to account for mean occupancies above 1. We incorporated an additional permissive state into the model, where two transcription factors can bind simultaneously, and fit a floating interaction energy between them (*ε*_*clash*_) (Figure 4D,E, Supplementary Figure 5, Supplementary Table 6). This 2-parameter (1 floating) model described the experimental data better than the exclusive model. Both the Akaike information criterion (AIC), and Bayesian information criterion (BIC) decreased significantly for the permissive model, which are measures of goodness-of-fit, and useful for conducting model selection when models differ in their number of parameters (penalizing the permissive model for its additional parameter, Supplemental Figure 5).

As the number of basepairs shared between binding sites increases, the mean occupancy decreases (as a function of concentration), and the probability of the 2 transcription factor-bound state is reduced, although it is still higher than initially expected at a large amount of overlap (3 basepairs, or 33% of the motif, Figure 4F). Furthermore, as we decrease site affinities, we observe a large decrease in occupancy in overlapping sites, suggesting that the transcription factor’s affinity for its binding site (*ε*_*bs*_) is important for compensating for the steric interference.

We further investigated binding to high-density clusters where individual sites differ in their affinities (Figure 4G, H). In this case, binding is dominated by the stronger binding sites in a given cluster. For the clusters of three mixed-affinity binding sites, the maximum occupancies and affinities were chiefly determined by the number of strong, non-overlapping binding sites present in the cluster.

### *In vivo* validation of low-affinity binding site cluster function in a synthetic gene regulatory system

Experimental methods and computational analyses designed to study gene regulation have traditionally been biased towards single sites of high-affinity or small clusters of high-affinity binding sites. Homotypic clusters of low-affinity sites are prevalent in eukaryotic regulatory sequences (Gotea et al. (2010)), yet the level of binding to these clusters in the *in vivo* context and the resulting impact on gene expression is uncertain. Having shown *in vitro* that at physiologically-relevant transcription factor concentrations an equal or greater mean transcription factor occupancy can be achieved by low-affinity clusters compared to an individual consensus site, we set out to investigate whether low-affinity site clusters are functional in the context of eukaryotic promoters *in vivo*.

Although our biochemical analysis showed that low-affinity binding site clusters are able to achieve high mean occupancies, several questions remained as to whether low-affinity binding site clusters would be functional *in vivo*. First, it is not a priori known whether high mean occupancies of transcription factors alone are sufficient to recruit the necessary regulatory machinery to a promoter and give rise to transcriptional activation. Consensus binding sites are able to achieve high mean occupancies of a single transcription factor, and this occupancy is further characterized by long-dwell times of the transcription factor on its binding site (Geertz et al. (2012)). Low-affinity clusters on the other hand achieve a similar mean occupancy in non-saturating conditions through a time-averaged occupation of several transcription factors bound to the cluster, and each transcription factor - DNA interaction exhibits a considerably shorter dwell time. It is a matter of debate whether high occupancies or long dwell times are necessary for *in vivo* function of regulatory regions. Second, we estimated that *in vivo* transcription factor concentrations may be high enough to give rise to sufficiently high occupancies of low-affinity clusters, but this assumption remained to be tested *in vivo*.

We first addressed the question of whether low-affinity binding site clusters can be functional *in vivo* by generating synthetic minimal promoters, driven by an engineered, exogenous Zif268 transcription factor. We inserted the 90 bp regulatory sequences that were characterized in our *in vitro* work approximately 200 bp upstream of the transcription start site in the minimal CYC1 promoter (Figure 5A). These sequences characterized *in vitro* were originally designed into a DNA backbone selected to prevent binding to known yeast transcription factors. We varied similar regulatory sequence parameters and maintained similar affinity-class definitions as for our low-affinity *in vitro* binding characterization.

**Figure 5:**
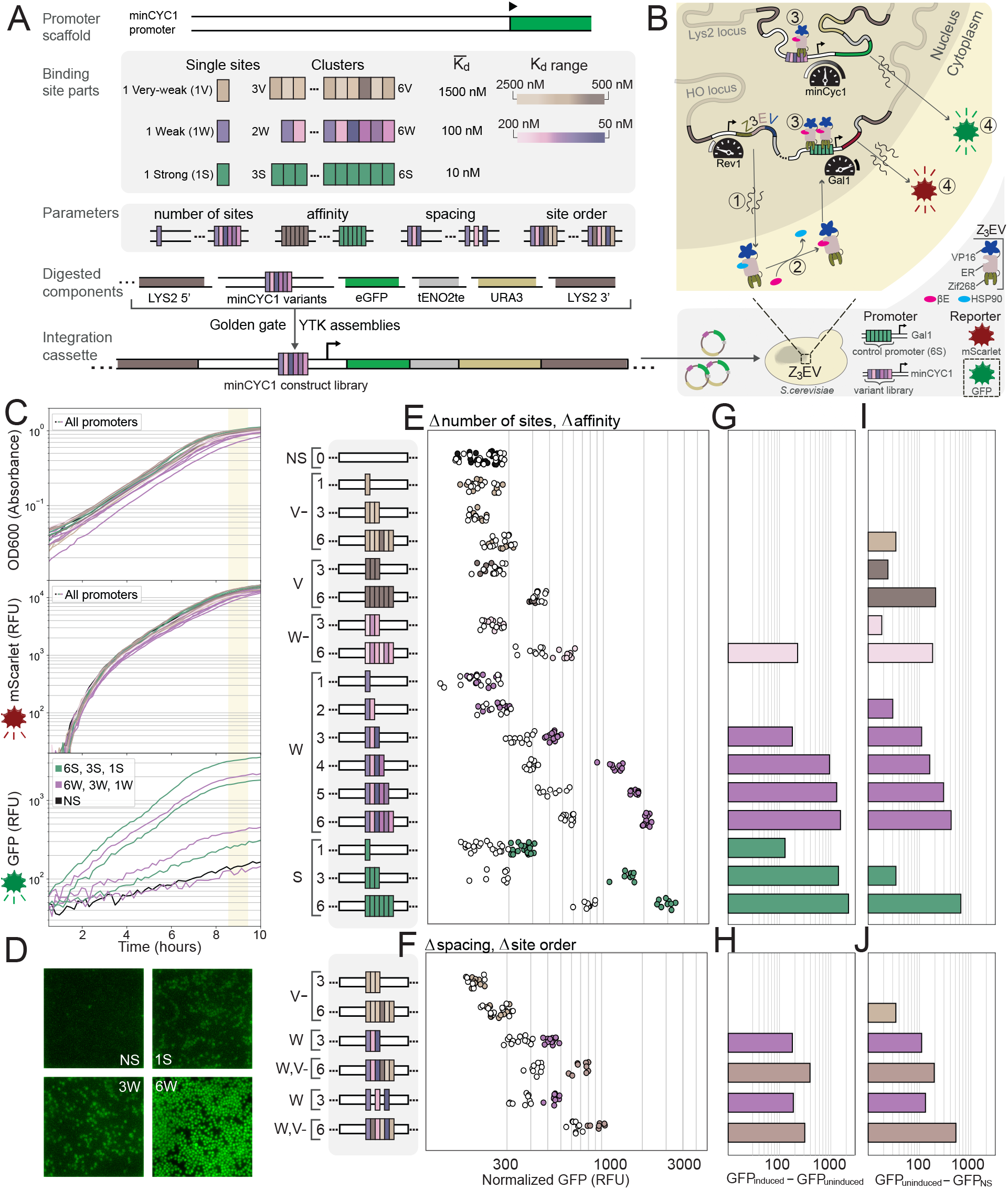
**(A)** Promoter legend. *In vitro* characterized Zif268 binding regions were inserted into a minCYC1 minimal promoter scaffold. Promoters were assembled into multi-cassette plasmids of standard parts driving GFP production, using the yeast-toolkit workflow (Lee et al. (2015)), and integrated into the LYS2 locus of a Z_3_EV parental strain (McIsaac et al. (2013)). **(B)** Z_3_EV is a synthetic transcription factor containing a Zif268 DNA-binding domain, an estrogen-responsive (ER) domain responsible for nuclear translocation upon *β*-estradiol induction and binding, and a VP16 activation domain. The parental strain was engineered to contain a synthetic GAL1 control promoter with 6 consensus binding sites driving mScarlet production, to quality-control Z_3_EV activity. **(C)** All strains displayed consistent growth (OD600, upper), and mScarlet production (middle). Strains varied in their GFP production depending on the minCYC1 promoter variant (bottom). Strains were characterized with single-cell fluorescence microscopy **(D)** and on a multi-mode platereader **(E-J). (E-F)** Strains were characterized either without (white, uninduced), or in the presence of saturating (200 nM) *β*-estradiol concentrations (colored, induced), over three independent experimental measurements. **(E)** The number of binding sites was varied from 1-6, and six distinct affinity classes were characterized (consensus binding sites (S), weak sites (W, 10X lower-affinity, W-, 20X lower-affinity), very-weak sites (V, 50X lower-affinity, V-, 100X lower-affinity), as well as a non-specific DNA target (NS). Promoters are organized from weakest (top) to strongest binding sites (bottom). **(F)** We varied the spacing between sites in clusters, characterized clusters with binding sites from mixed affinity classes, and changed the ordering of binding sites in mixed-affinity clusters. **(G-H)** A Z_3_EV-specific response was quantified as the difference in fluorescence intensity between induced and uninduced conditions. **(I-J)** A non-specific response was quantified as the difference in fluorescence intensity in uninduced conditions between a given promoter strain and the NS promoter-containing strain. The non-specific response also showed strong correlation with gene expression measurements of the same promoter library integrated in a BY4741 yeast strain lacking Z_3_EV (Supplementary Figure 6).

We used the yeast toolkit (YTK) 2.0 workflow (Lee et al. (2015)) to assemble integration cassettes with our engineered minCYC1 promoters upstream of GFP. We integrated these synthetic cassettes into the Lys2 locus of a modified Z_3_EV-containing parental yeast strain (Figure 5B) (McIsaac et al. (2013)). Z_3_EV is a synthetic transcriptional activator with three domains: the Zif268 DNA-binding domain, an estrogen receptor element that drives nuclear translocation in the presence of *β*-estradiol, and a VP16 domain that recruits transcriptional machinery to activate transcription. We also included a control promoter into the parental strain which drives mScarlet expression under the regulation of 6 consensus Zif268 binding sites in a synthetic GAL1 promoter backbone. The same parental strain was used for all strains generated, allowing us to validate the correct functioning of the inducible gene expression system to ensure consistency across strains.

Yeast strains were either induced with a high concentration of *β*-estradiol (200 nM), or not uninduced (0 nM), and measured in the mScarlet and GFP channels on a multi-mode platereader (Figure 5C) and cross-validated by direct fluorescence microscopy imaging (Figure 5D, Supplementary Figure 6). All strains exhibited highly consistent growth rates and mScarlet expression from the control-promoter (Figure 5C, Supplementary Figure 6E-H). In contrast, the cluster variants in our engineered promoter library led to a wide range of gene expression levels (GFP) that depended on cluster configuration and binding site affinities. We normalized GFP intensities by cell densities, and averaged this normalized gene expression signal for all readings between 8.5 hours and 9.5 hours after induction (Figure 5E, F). Each datapoint in Figure 5E corresponds to an independent replicate, where generally around three replicates were characterized in each experiment, and the experiment was repeated from three to six times for each strain.

All promoters containing 1, 3, or 6 consensus binding sites gave rise to functional activation and expression of GFP. The minCYC1 promoter containing a single consensus binding site achieved only modest expression levels, whereas 3 consensus binding sites led to very high expression of GFP (Figure 5E-H). Increasing the number of consensus binding sites from 3 to 6 led to slightly higher gene expression levels, potentially indicating that expression levels are near the upper limit of what the minCYC1 promoter can support. The output of a promoter containing no binding sites (NS) on the other hand remained low, both in the induced and uninduced conditions. We tested binding site clusters containing between 1 and 6 very weak binding sites with average *K*_*D*_ values 2 orders above the *K*_*D*_ of the consensus sequence. None of these promoters led to a measureable increase in gene expression when comparing the induced to the non-induced state, with the possible exception of a promoter containing 6 binding sites with slightly higher binding affinities (W-6). We started observing a difference in gene expression between the induced and non-induced conditions, when testing clusters of weak binding sites with average *K*_*D*_ values one order of magnitude above the *K*_*D*_ of the consensus sequence. Clusters of as few as 3 weak binding sites (W3) surpassed the single consensus binding site in its ability to activate gene expression, while clusters of 4, 5, or 6 weak binding sites achieved a similar level of gene expression compared to clusters composed of three consensus binding sites. Therefore, we were able to show in this minimal synthetic gene regulatory system, that small clusters of low-affinity binding sites can drive gene expression equivalently to clusters of consensus binding sites.

Interestingly, we found that not only did promoters vary in their induced levels of GFP-expression, but they also varied significantly in uninduced gene expression in a way that depended consistently on the promoter class. Increasing numbers of binding sites within a cluster gave rise to increasing levels of leakiness in the uninduced condition (Figure 5I-J). The highest level of leakiness was observed for a cluster of 6 consensus sequences (S6), and the level of leakiness scaled with the number of weak binding sites in a cluster (W1-6). Even non-functional clusters of very weak binding sites gave rise to increased leakiness (V6 and W-6). The one exception to this general observation was the promoter containing 3 consensus sequences (S3) which achieved high levels of induction without any significant increase in leakiness. We reasoned that this response could either be specific and due to Z_3_EV transcription-factor leak under non-inducing conditions, or non-specific, resulting from the binding of alternative endogenous transcription factors. Therefore, we characterized the entire promoter library in a wildtype BY4741 yeast strain lacking Z_3_EV, and discovered that the level of expression correlated strongly with uninduced gene expression in the Z_3_EV parental strain (R^2^ of 0.9, Supplementary Figure 6). This confirmed that the leakiness in the uninduced state was due to non-specific binding of one or more endogenous transcriptional activators.

Our library of well-characterized regulatory sequences where mean occupancy and affinity vary independently is useful for trying to understand which binding properties predominantly drive transcription from clusters. In eukaryotes, there are limited and conflicting examples where gene expression has been explained using a reduced set of interpretable binding properties (Estrada et al. (2016); Lickwar et al. (2012); de Boer et al. (2020); Segal et al. (2008); Rajkumar et al. (2013); Mogno et al. (2013)). We used our *in vitro*-characterization data to predict the mean occupancies of transcription factor molecules binding to clusters in the Z_3_EV promoter library *in vivo*. We compared these predictions to the TF-specific promoter responses (Figure 6).

**Figure 6:**
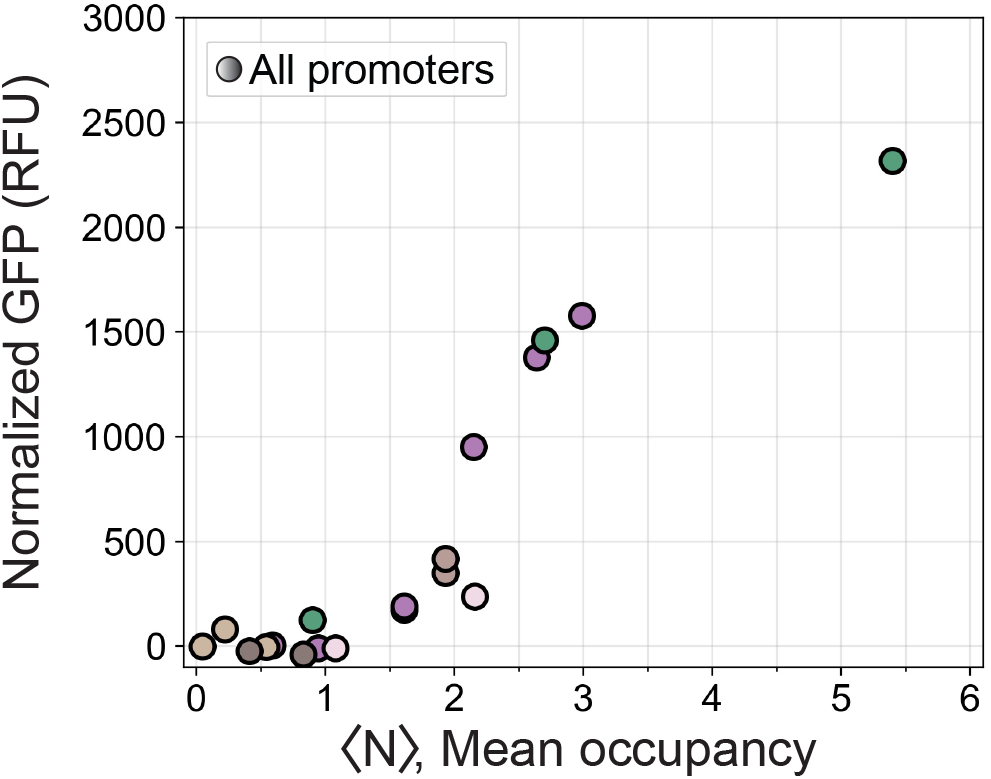
Thresholded relationship of gene expression (GFP normalized by OD600) as a function of predicted mean occupancy (based on *in vitro* characterization data), for the full Z_3_EV strain library. Color scheme is the same as in Figure 5.

Our model suggests that the minCYC1 promoter regulates gene expression through a simple thresholded relationship with the average number of bound transcription factor molecules, where up until an average of around 1 transcription factor bound to the promoter, gene expression is largely silent, and after this threshold, expression depends primarily on the average number of transcription factor molecules bound rather than on their affinity for their binding sites. For instance, a cluster composed of 5 weak binding sites (W5) or 3 strong binding sites (S3) differ in affinity by an order of magnitude, but show similar levels of gene expression which can be explained by their predicted mean occupancies. These results do not vary qualitatively within a reasonable range of expected parameter values (Supplemental Figure 6). Thresholding can also be observed qualitatively, since one weak site (W1), or two weak sites (W2) show no impact on expression, but each additional weak site from then on increases expression significantly (Figure 5E). Furthermore, adding 3 very-weak sites to a very-weak cluster (moving from V-3 to V-6) does not increase transcription factor-specific expression, while appending these same 3 very-weak sites just after a cluster that has a higher starting occupancy (moving from W3 to W3V-3) causes a transcription factor-specific increase in promoter output (Figure 5F). We observed no difference in expression level when we changed spacing between binding sites (W3 compared to W’3), and little to no difference when we changed the ordering of sites in promoters containing binding sites from different affinity classes (W3V-3 compared to W3V-’3V).

### *In vivo* validation of low-affinity binding site cluster function in a native gene regulatory system

Having established that clusters of low-affinity binding sites are functional in a synthetic gene regulatory network, we next addressed the question of whether low-affinity binding sites clusters would also be functional in a native promoter regulated by a transcription factor expressed at physiological concentrations. We turned to the well-studied inorganic phosphate regulatory network in *S. cerevisiae* to address this question. The PHO5 promoter is one of the best understood promoters of the inorganic phosphate regulatory network in yeast. It is activated in response to phosphate starvation, which causes the master regulator Pho4 to localize to the nucleus leading to binding to a Pho4 binding site and an overlapping Pho2 site located in a nucleosome-free region (NFR) (Figure 7A). This binding results in the displacement of a nearby nucleosome (in the -2 position), which in turn makes a higher-affinity Pho4 site and two nearby Pho2 binding sites available for binding which are nominally in a nucleosome-occluded region (NOR). This mechanism is believed to decouple the promoter’s activation threshold from its dynamic range, where as a first approximation the Pho4 and Pho2 binding sites in the NFR are considered to confer the promoter with its threshold and all binding sites in the NFR and NOR determine gene expression levels after that threshold is met (Lam et al. (2008); Rajkumar et al. (2013)).

**Figure 7:**
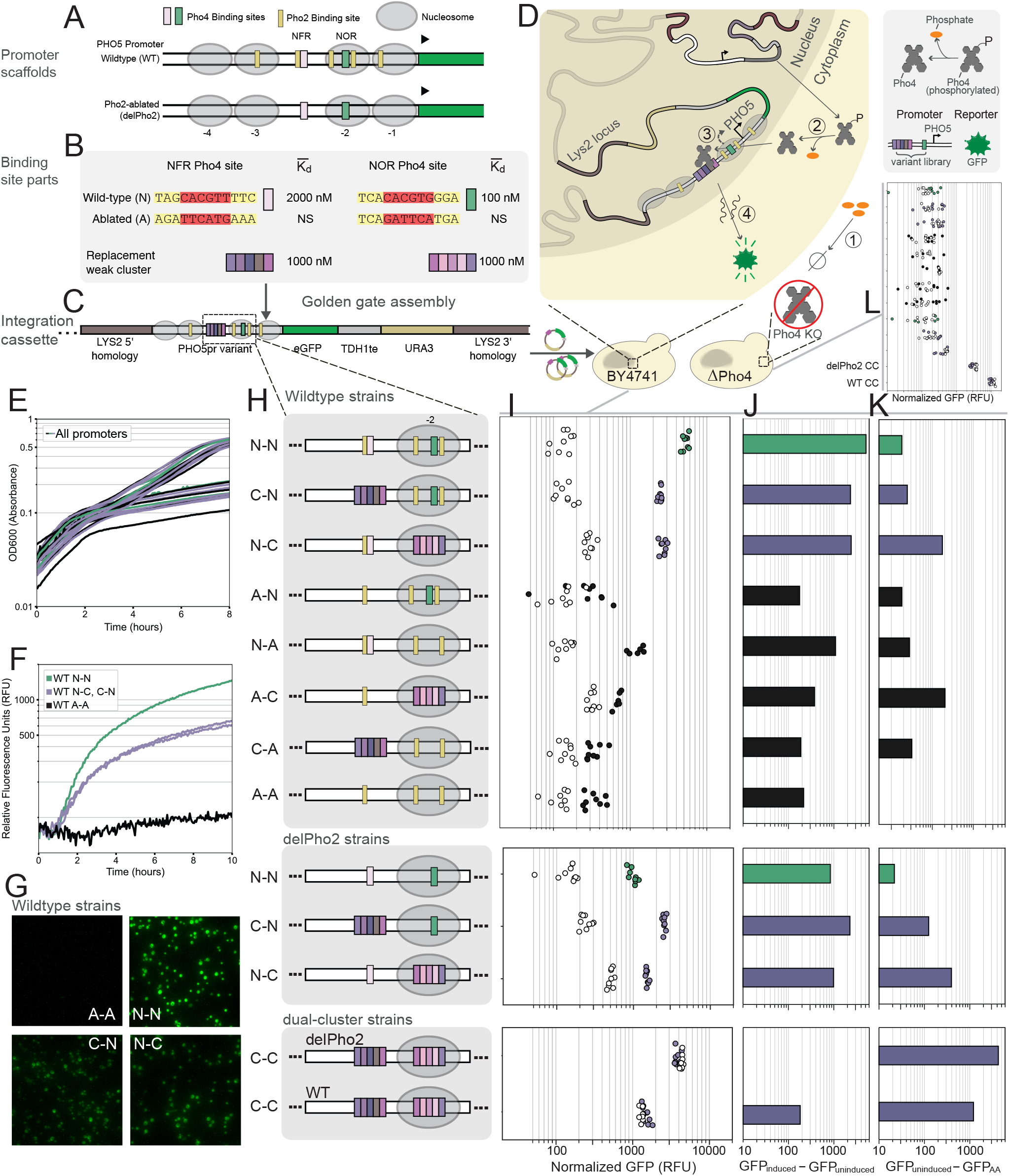
**(A)** PHO5 promoter scaffolds in which native Pho4 binding sites are replaced with low-affinity Pho4 clusters. **(B)** Legend of Pho4 binding site parts. Native binding regions containing Pho4 and Pho2 sites either have their Pho4 binding sites ablated, or are replaced with clusters of low-affinity Pho4 binding sites. **(C)** Promoters were assembled into a multi-cassette plasmid and integrated into the LYS2 locus of either BY4741 or Pho4 knockout (*δ*Pho4) yeast strains. **(D)** Schematic of the response (induction) to phosphate starvation. **(E)** OD600 time series measurements for the full BY4741 library of promoters, both for uninduced and induced conditions. **(F)** Induction time series measurements for select cluster-containing strains. **(G)** Strains were also characterized by single-cell fluorescence microscopy. **(H-J)** A Pho4-specific response was quantified as the difference in fluorescence intensity between induced and uninduced conditions. **(K)** A non-specific response was quantified as the difference in fluorescence intensity in uninduced conditions between a given promoter strain and the ablated-ablated (AA) promoter strain. **(L)** The non-specific response can also be seen in a Pho4 knockout strain, with a similar relationship across promoter variants (Supplementary Figure 7).

Significant effort has been invested to uncover the functional relevance of the individual binding sites in the PHO5 promoter. A Pho2 binding site in the NFR has been found to recruit chromatin remodellers and to promote cooperative binding of Pho4, while the Pho2 sites in the NOR have been identified as necessary for efficient activation by Pho4 (Barbaric et al. (1998, 1996)). Ablation of Pho2 binding sites has been found to decrease gene expression up to 10-fold (Rajkumar et al. (2013); Barbaric et al. (1998)), and a Pho2 knockout strain exhibits close to no PHO5 promoter expression compared to a wildtype strain (Fascher et al. (1990)). However, Pho4 overexpression has also been shown to compensate partially for the loss of expression due to Pho2 binding site ablation (Barbaric et al. (1998)), or even for the loss of expression in Pho2 knockout strains (Fascher et al. (1990); Nourani et al. (2004); Korber and Barbaric (2014)), presumably through an increase in occupancy of the native Pho4 binding sites. Based on this picture, and since weak clusters of Pho4 binding sites attain similar to greater occupancies than the individual consensus or native sites at relevant transcription factor concentrations *in vitro* (Figure 2), we hypothesized that weak binding site clusters might achieve sufficiently high Pho4 occupancies in the PHO5 promoter and reproduce part of the functionality conferred by the native regulatory sequence, both in the NFR and NOR.

We replaced the native Pho4 binding sites together with their neighboring Pho2 sites, in the NFR or NOR, with clusters of 5 low-affinity Pho4 binding sites (weak or very-weak) (Figure 7A-C) that we had characterized *in vitro* (Figure 2). We conducted this replacement either in a native promoter scaffold (WT), or a scaffold entirely depleted of known Pho2 binding sites (delPho2), in order to decouple the Pho2-specific effect from the effect of Pho4 occupancy. PHO5 promoters were assembled into a cassette driving GFP reporter expression (Figure 7C), and integrated into the LYS2 locus of either a wildtype yeast strain (BY4741) or a Pho4 knockout strain lacking Pho4 (Figure 7D). All strains were sequence-verified, and exhibited consistent growth irrespective of their PHO5 promoter variant (Figure 7E). The positioning of nucleosomes in the PHO5 promoter has been characterized extensively and found to be robust to promoter context, such that it remains consistent when the promoter is located in an alternate locus (e.g. the LYS2 locus) and for a large degree of modification to the promoter sequence (Korber and Barbaric (2014); Rajkumar et al. (2013)).

Our results on deleting native Pho4 binding sites in the NFR and NOR regions are in concordance with previous characterizations of the PHO5 promoter (Lam et al. (2008); Rajkumar et al. (2013)) (Figure 7E-J, Supplementary Figure 7). Indeed, we observed greater expression for the WT promoter with a native NFR but an ablated NOR Pho4 site (WT N-A), compared to the promoter where both NFR and NOR Pho4 sites were ablated (WT A-A). This level of expression is then scaled with the addition of the native NOR Pho4 site (WT N-N). On the other hand, the promoter with an ablated Pho4 site in the NFR and a native Pho4 site in the NOR (WT A-N), resulted in similar expression to the ablated-ablated promoter (A-A), presumably because the nucleosome was not displaced effectively upon induction.

Compared to the wildtype A-N promoter, replacing the Pho4 binding site in the NFR (which also deleted the Pho2 binding site) with a cluster of low-affinity Pho4 binding sites (WT C-N) recovered Pho4-specific expression. Furthermore, compared to the N-A promoter, replacing the NOR (which also deleted the Pho2 binding sites) with a cluster of low-affinity sites (WT N-C) also improved expression compared to the N-A promoter. Promoters with no native Pho4 binding sites and a low-affinity binding site cluster in the NFR (C-A) or NOR (A-C) lost function similar to promoters with the corresponding native Pho4 binding sites present (N-A and A-N, respectively). The wildtype C-N promoter showed significantly higher expression than wildtype C-A promoter (and A-N showed ablated levels of expression), suggesting that the low-affinity cluster in the NFR may contribute to nucleosome displacement to uncover the NOR. Our results show that low-affinity clusters can generate similar expression levels of the PHO5 promoter compared to what is conferred by the native cis-regulatory sequence of the NFR and the NOR. Partial recovery of function was expected based on the aforementioned work demonstrating that overexpressing Pho4 (presumably increasing its concentration and occupancy) will only partially compensate for Pho2 site ablation (Barbaric et al. (1998); Fascher et al. (1990)).

Indeed, when all Pho2 binding sites are deleted from the promoter, gene expression from the native promoter (delPho2 N-N) drops by around 6-fold, consistent with previous reports (Barbaric et al. (1998); Rajkumar et al. (2013)). Remarkably, in this Pho2-depleted scaffold, replacing the native Pho4 sites in either the NFR or NOR with clusters of low-affinity binding sites (delPho2 C-N, or delPho2 N-C, respectively) actually resulted in a similar or greater level of gene expression than what was achieved by the native binding sites lacking the neighboring Pho2 sites. This implies that the Pho4-specific impact on gene expression of low-affinity Pho4 clusters is equal to or greater than for the native Pho4 binding sites and its potential co-operative interaction with Pho2. This then suggests that low-affinity clusters attain significant occupancy, which we previously established *in vitro* and in a minimal promoter *in vivo*, and can be extended to native and more complex promoter settings.

Expression in the delPho2 C-N strain does not decrease compared to the WT C-N strain (containing all Pho2 binding sites except for the one replaced by the cluster in the NFR), whereas compared to the WT N-C the delPho2 N-C strain shows lower expression, suggesting that the Pho2 binding site in the NFR is of greater importance to gene expression as has been suggested in the literature (Barbaric et al. (1998)). Indeed, our results suggest that ablating the nucleosome-free Pho2 site hinders the efficacy of the NFR, likely in part by reducing cooperative binding to Pho4 (Barbaric et al. (1998)). However, our results suggest that the function of this regulatory region can be compensated for by strengthening the region’s Pho4 mean occupancy through a low-affinity Pho4 binding cluster, leading to a higher level of gene expression.

Interestingly, similar to the Z_3_EV promoter system, non-specific leaky expression correlated with the number of low-affinity binding sites (Figure 7K). Dual-cluster replacement resulted in significant non-specific expression, which did not depend on Pho4. This is evidenced by the fact that expression did not depend on induction in the wildtype background. Furthermore, these strains were the only ones to exhibit a high level of expression at timepoint zero of induction, and their expression did not increase over time. This was the case for dual-cluster replacement both in the native promoter scaffold and in the scaffold depleted of Pho2 binding sites. In further agreement, these promoters showed significant expression in the *δ*Pho4 background (Figure 7L). Replacement of the NOR alone by a weak-cluster was enough to drive a moderate level of non-specific expression, while this was not the case for cluster-replacement in the NFR. Aside from the above discussed strains, all other strains showed little to no non-specific expression. They exhibited consistent, ablated levels of gene expression in the absence of induction, and in the *δ*Pho4 background.

## Discussion

Our current understanding of how transcription factors bind to genomic regulatory regions to drive transcription has developed largely through experiments that characterize binding to individual binding sites, often with a focus on high-affinity consensus sites. The functional relevance of low-affinity binding sites, particularly clusters of low-affinity binding sites is less-well understood (Kribelbauer et al. (2019)). Methodological challenges associated with measuring binding of multiple transcription factors to low-affinity clusters has prevented precise measurements of how binding to these clusters compares with binding to individual consensus binding sites, creating uncertainty about the significance of low-affinity clusters. Indeed, through prior state-of-the-art biochemical methods used to measure collective binding in high-throughput, occupancy appeared driven primarily by strong binding sites (Levo et al. (2015)). Reports in the literature identified *in vivo* examples where low-affinity clusters are functionally important (Crocker et al. (2015); Kribelbauer et al. (2019); Crocker et al. (2016); Gaudet and Mango (2002); Farley et al. (2015); Jiang and Levine (1993), or exhibit properties like binding synergy (Segal et al. (2008); Castellanos et al. (2020)). But, the lack of a systematic analysis *in vitro* or *in vivo* limited us to inductive reasoning when trying to apply these findings to new contexts. By combining *in vitro* biochemical analysis with *in vivo* functional studies of gene regulation in the eukaryotic model system *S. cerevisiae* we were able to provide insights into how transcription factors interact with low-affinity clusters and were able to show that these clusters are functional *in vivo*.

We applied a high-throughput *in vitro* method to characterizing collective transcription factor binding to DNA sequences consisting of arrays of binding sites ranging over orders of magnitude in affinity. This permitted us to develop a quantitative biophysical understanding of how transcription factors interact with binding site clusters, particularly to clusters consisting of low-affinity binding sites. Clusters of low-affinity binding sites achieved similar levels of occupancy as single high-affinity binding sites and we predicted that this level of occupancy can be achieved at physiologically relevant transcription factor concentrations. We verified this effect with two distinct transcription factors representing two of the largest transcription factor structural classes: zinc finger and bHLH. Small clusters of 2-6 binding sites with individual binding site affinities one order of magnitude lower than the consensus sequence were able to achieve this effect. We furthermore challenged the notion that overlapping binding sites are mutually exclusive due to steric effects, and thus would allow only one transcription factor to bind to overlapping sites at any given time. In fact, clusters of strong binding sites with a one bp overlap of a 9 bp consensus sequence were not mutually exclusive and were simultaneously bound by several transcription factors. Even a 3 bp overlap of a 9bp consensus motif permitted some co-occurring binding, although here the steric factor strongly dominated. Similar trends were observed for weak binding site clusters with 1 and 3 bp overlapping sequences, indicating that strong binding sites permit and support partial binding of a transcription factor, allowing full occupancy of overlapping binding sites.

The high levels of occupancy achieved by low-affinity binding site clusters as determined by our *in vitro* characterization suggested that low-affinity binding site clusters may indeed be functional *in vivo*. We therefore first tested low-affinity binding site clusters in an engineered synthetic system *in vivo*. We placed low-affinity Zif268 binding site clusters in the minimal CYC1 promoter and used a *β*-estradiol inducible Zif268 transcription factor (Z_3_EV) to test whether these promoters could give rise to expression. We found that binding site clusters of 3 to 6 weak binding sites with individual affinities one order of magnitude below the consensus sequence were able to activate transcription to similar levels as a promoter containing 3 consensus sequence binding sites. Therefore, in this synthetic system, small clusters of low-affinity binding sites are functional. We also observed a concurrent increase in promoter leakiness which we characterized and attributed to non-specific interactions of an endogenous transcription factor binding to these binding site clusters. Although clusters of low-affinity binding sites were functional in the synthetic Z_3_EV-minCYC1 system, it was possible that this was due to high, super-physiological expression levels of the Z_3_EV transcription factor. To control for this uncertainty we tested whether low-affinity binding site clusters could substitute single high-affinity binding sites in a native gene regulatory network. We substituted native binding sites in the nucleosome-free and nucleosome-occluded regions of the well-characterized PHO5 promoter, which is regulated by endogenous levels of the master regulator Pho4. In this context low-affinity binding site clusters were also able to functionally replace single high-affinity binding sites, recovering a large proportion of the native PHO5 gene expression level. This was particularly impressive considering that substitution of the native Pho4 binding sites in these two locations also ablated the neighboring Pho2 binding sites, which are known to further strengthen the effect of Pho4 binding to these single target sites and contribute to transcriptional activation of this promoter.

These findings have several important implications on our current understanding of transcriptional regulation, computational methods for predicting gene regulatory function, gene regulatory network evolution, and engineering of synthetic gene regulatory networks. We have shown that low-affinity binding site clusters are effective at activating transcription *in vivo* suggesting that for computational approaches to gene regulatory function prediction and transcription factor binding prediction based on ChIP-seq data, that low-affinity binding sites warrant closer consideration, and that clusters of low-affinity binding sites can be highly functional.

On a more fundamental level, it was not known whether transcription factor dwell time or occupancy is the critical parameter that determines whether a binding region can give rise to functional gene expression. A single high-affinity binding site and clusters of low-affinity binding sites can both give rise to high occupancies at physiologically relevant transcription factor concentrations. But how this occupancy is achieved is fundamentally different between these two cases, with a single high-affinity binding site achieving high occupancy due to long transcription factor dwell times (low off-rates), whereas low-affinity binding site clusters achieve high occupancy as an ensemble average of many interactions of short dwell times (high off-rates). Our findings show that both systems can give rise to transcriptional activation and suggests that long transcription factor dwell times are not required for transcriptional activation. This may not apply to transcriptional repression where it is possible that long dwell times are critical for efficient repression. It will be interesting to assess whether low-affinity binding site clusters can function as effective repressive regulatory regions or not. Finally, whether there is an observable difference in regards to gene expression noise between low-affinity binding site clusters and single high affinity binding sites would also warrant further exploration (Azpeitia and Wagner (2020)).

From an evolutionary perspective, low-affinity binding site clusters are of considerable interest as they could serve as a functional intermediate between a non-regulated promoter and a highly evolved promoter containing one or more consensus sequence binding sites. It is interesting that clusters of low-affinity binding sites are functional, but that they also seem to come with an additional cost which is non-specific regulation or cross-talk. Small clusters of low-affinity binding sites thus might be easier to form from a non-specific sequence background, but are then under continued pressure to further evolve to more specific high-affinity binding sites. It is striking that as few as 3 low-affinity binding sites are sufficient to generate a highly functional regulatory region. Likewise, when approaching this notion from a transcription factor centric perspective, there exist many more possible low-affinity binding sites for any given transcription factor than specific sites, which also enables regulatory regions to evolve more readily but also is the reason for non-specific interactions and cross-talk. Transcriptional regulation has traditionally been considered as being encoded in strong, well-defined binding sites. Results from de Boer et. al. supported a more finer-grained view of transcription by demonstrating that gene expression in promoters of *S. cerevisiae* is largely driven through the collective action of many weak regulatory contributions (de Boer et al. (2020); Tanay (2006)). Holding total occupancy constant, many weaker sites may allow for a greater number of binding sites and different transcription factors (Tanay (2006)), increasing the potential of network connectedness, or the average degree of nodes found in a eukaryotic gene regulatory network.

Synthetic promoters to date have predominantly made use of consensus binding sites, high-affinity binding sites, clusters thereof (Khalil et al. (2012)), and most recently clusters of lower affinity binding sites in combination with transcription factor cooperativity (Bashor et al. (2019)). As engineering synthetic promoters and entire gene regulatory networks matures it is likely that increasingly precise gene expression levels will become necessary. This to some extent may be achievable with relatively few single binding sites of various affinity, but it may be worthwhile considering the use of low-affinity binding site clusters. Low-affinity binding site clusters may be more resilient to mutations for example. First, a single deleterious mutation in a cluster of several low-affinity binding sites will have a much smaller impact on the resulting gene expression level as a mutation in a single high affinity binding site. Second, a single base change in a single high-affinity binding site will result in a large decrease in binding affinity, whereas a single base change in a low-affinity site will lead to a much smaller relative decrease in affinity, or may even be silent in regards to affinity.

In summary we conducted a systematic characterization of low-affinity binding site clusters by conducting a quantitative biochemical analysis *in vitro* as well as functional studies *in vivo*. This work provided insights into how transcription factors bind to such clusters and established that a small number of low-affinity binding sites in a local cluster can be highly functional *in vivo*. These insights improve our current understanding of gene regulatory networks, and our ability to engineer sophisticated gene regulatory networks *de novo*.

## Acknowledgments

S.J.M. and I.I. thank Prof. David Shore (University of Geneva) for many insightful discussions. This work was supported by the following grants to S.J.M.: a Swiss National Science Foundation SystemsX.ch IPhD grant (SNF:51PHP0 157292 / SysX:2014/242), a Swiss National Science Foundation Sinergia grant (189910), and by the European Research Council under the European Union’s Horizon 2020 research and innovation program grant 723106.

## Author contributions

A.S. performed all *in vitro* experiments, data analysis and modeling used in this manuscript. M.L. performed all *in vivo* experiments, M.L. and A.S. performed data analysis. I.I. and S.J.M. designed the initial iMITOMI method which was further refined by A.S. I.I. and S.J.M. designed the Pho4 target library. A.S. and S.J.M. designed the Zif268 target library. I.I. designed the recombinant proteins and cloned the expression plasmids. I.I. and A.S. generated the recombinant proteins. I.I. collected and analyzed preliminary Pho4 data. E.O. supported I.I. and A.S. with development of the iMITOMI method, modeling, and data analysis. S.C., M.L., and A.S. generated the yeast strains. A.S. and S.J.M. wrote the manuscript with input from all authors. S.J.M. conceived and directed the study.

