## Supplementary Information for "Systematic analysis of low-affinity transcription factor binding site clusters *in vitro* and *in vivo* establishes their functional relevance"

#### Part I

### Supplementary figures and tables

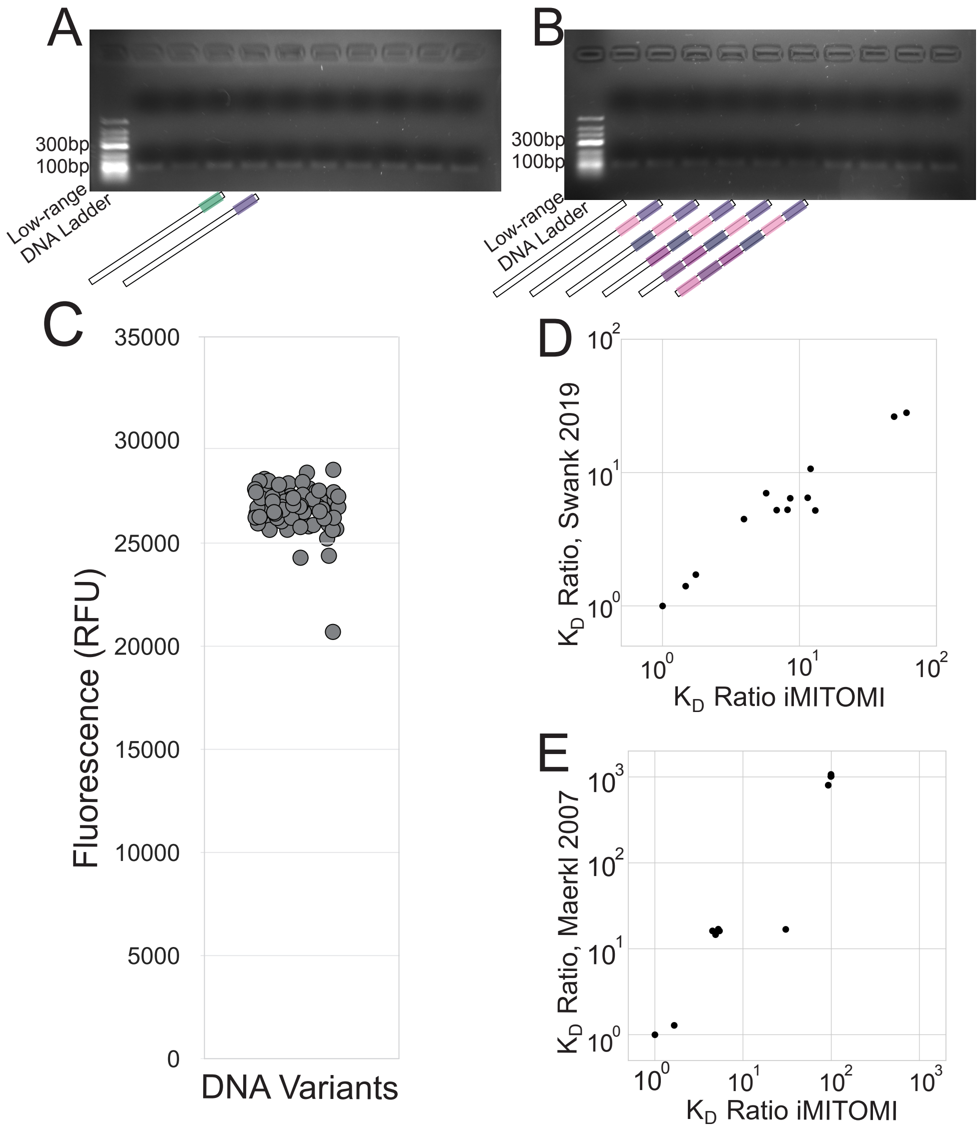

Figure S1: **(A, B)** Agarose gels for iMITOMI 90 bp PCR products (see Section 1) corresponding to the multi-weak Zif268 DNA target library characterized in (Figure 2C). **(C)** Fluorescence measurements of concentration-equalized iMITOMI DNA targets in a spotting plate before spotting. **(D, E)** Similar affinities and specificities were obtained when single site DNA targets were characterized on iMITOMI for binding to Zif268, and Pho4, as compared to Swank et al. (2019) and Maerkl and Quake (2007a), respectively. The $K_D$ Ratio represents the $K_D$ divided by the $K_D$ of the strongest binding site analyzed in this study.

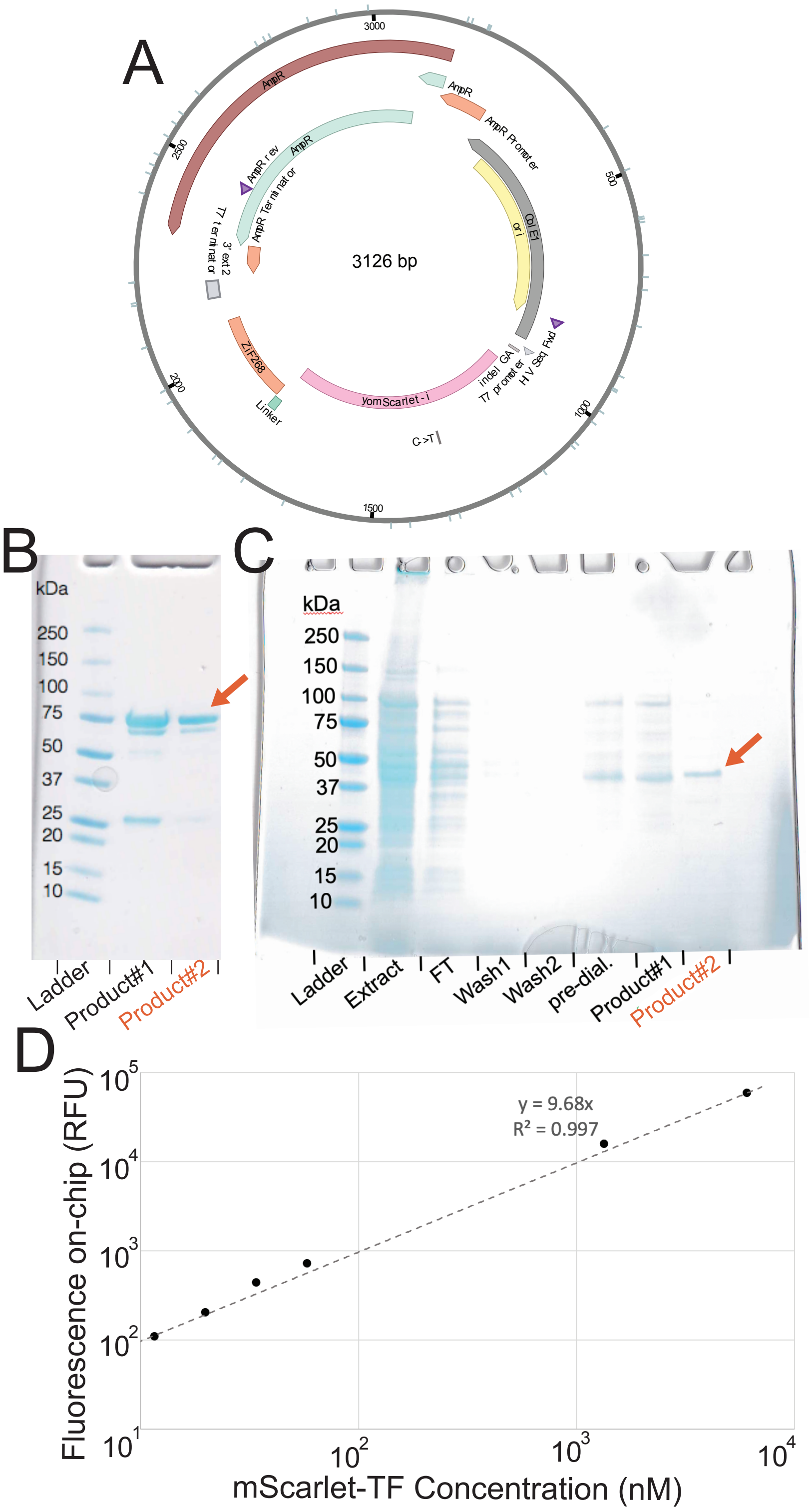

Figure S2: Characterization and calibration for purified proteins. Caption on the next page.

Figure S2: **(A)** Plasmid for 5'-mScarlet-Zif268-6xHis-3' construct under a T7 promoter. A similar construct was used for the Pho4 transcription factor. **(B, C)** Proteins were his-tag purified and ran on a denaturing PAGE gel for Pho4 (**B**) and Zif268 **(C)**. **(B)** Compared to Product #1, Product #2 was additionally buffer exchanged in an Amicon spin-column. **(C)** Product #2 is from an entirely different purification batch. In both cases the higher purity Product #2 was used for all iMITOMI experiments in this study. **(D)** Calibration curve relating fluorescence measurements on-chip to mScarlet-transcription factor concentration quantitated by a Bradford assay.

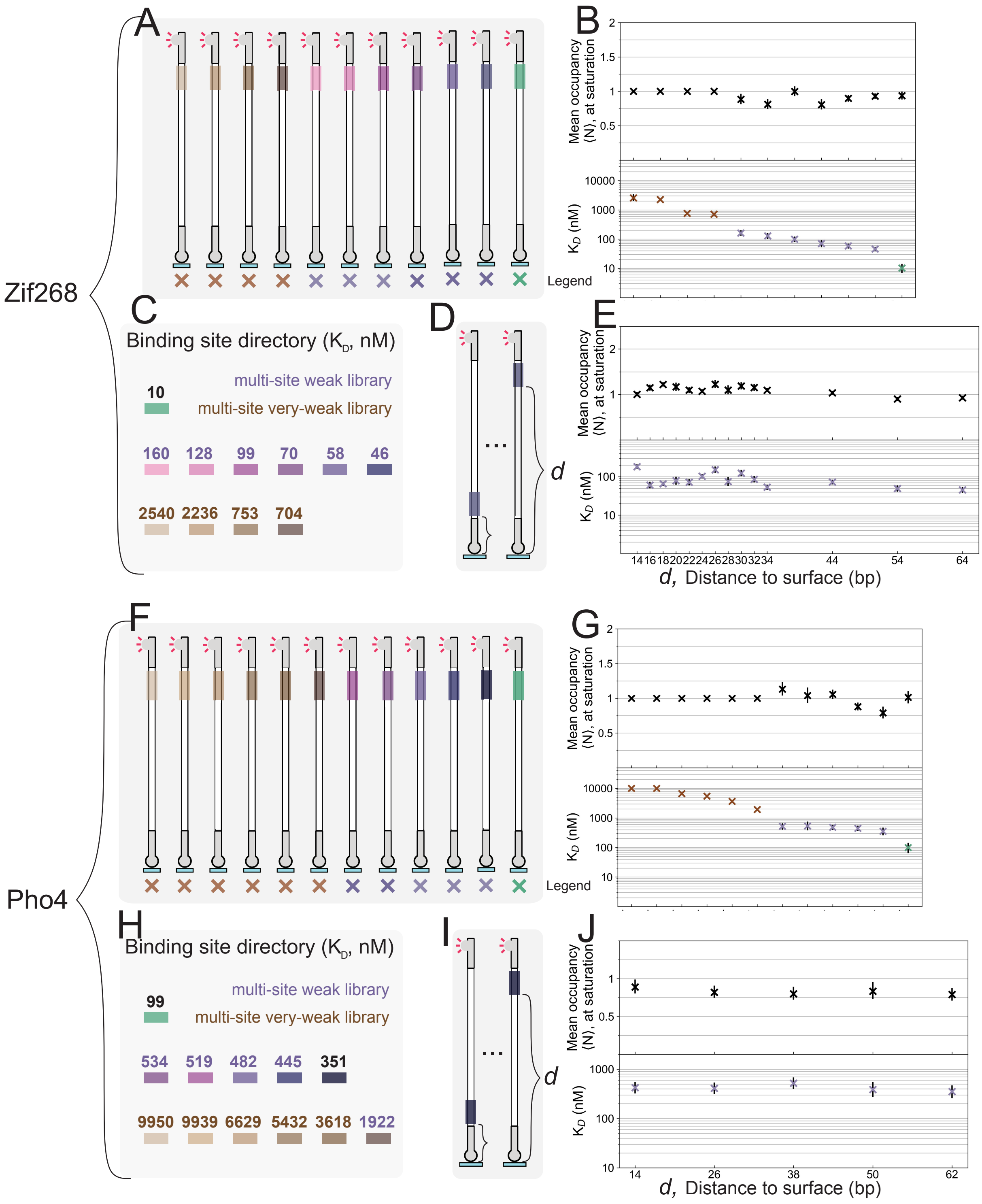

Figure S3: Binding site directories and single site positions scans.

Caption on the next page.

Figure S3: **(A)** Single site DNA targets for binding sites used to build Zif268 clusters throughout the study. **(B)** Mean occupancies at saturation (upper) and $K_D$ values (lower) for targets in **(A)**, in the same order from left to right. For single weak binding sites, the $\langle N \rangle_{\max}$ parameter was fixed. **(C)** Color-coded Zif268 binding site directory with affinity values in nanomolar. **(D)** DNA targets corresponding to the position scan in **(E)**, where a Zif268 weak binding site's distance ($d$) from the chip surface was varied across targets in the library. **(E)** Characterization data for the Zif268 single site position scan library. **(F)** Single site DNA targets for binding sites used to build Pho4 clusters throughout the study. **(G)** Mean occupancies at saturation and $K_D$ values for targets in **(F)**. **(H)** Color-coded Pho4 binding site directory with affinity values in nanomolar. **(I)** DNA targets corresponding to a Pho4 binding site position scan. **(J)** Characterization data for the Pho4 position scan library. Similarly to **(E)**, the distance to the chip's surface did not significantly impact binding.

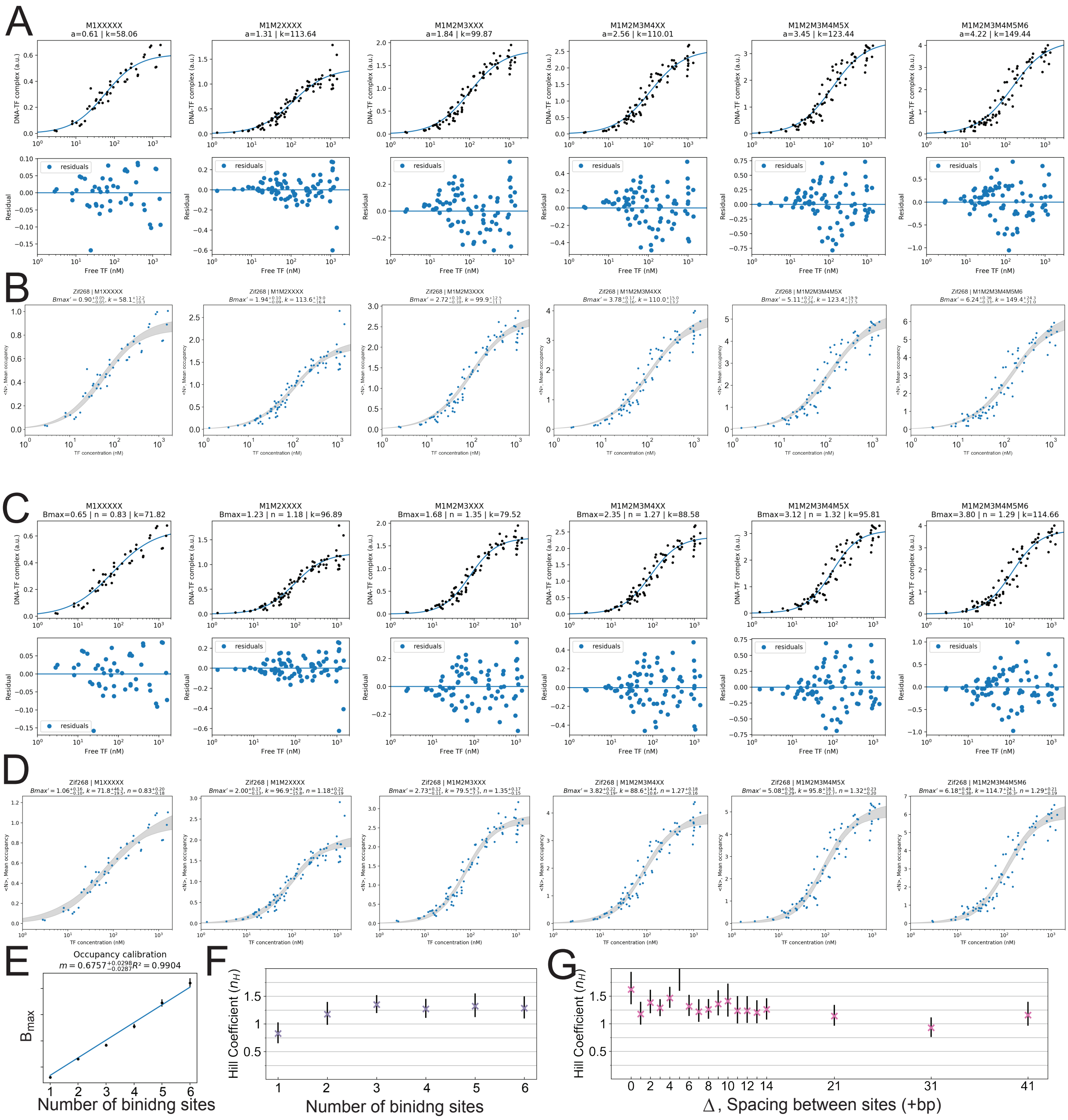

Figure S4: Data and model fits (SBC, Hill) for the Zif268 multi-weak library in Figure 2. Caption on the next page.

Figure S4: **(A)** Data was fit with a 2-parameter saturation binding curve (upper) (Equation 7). Residuals for the fit (lower). **(B)** Parameter space was explored using Markov chain Monte Carlo (MCMC) to obtain a 5 to 95% confidence interval range. Data is normalized by the step increase in $B_{max}$ (Equation 9). **(C, D)** Similar plots to **(A, B)** instead using a Hill model (8). **(E)** Step increase in $B_{max}$ with additional binding sites for the saturation binding model fits in **(A)**. **(F, G)** Hill coefficients ($n_H$) for the multi-site weak library in **(C, D)** and for the two-site gap-scan library targets (Figure 3) fit with a Hill model.

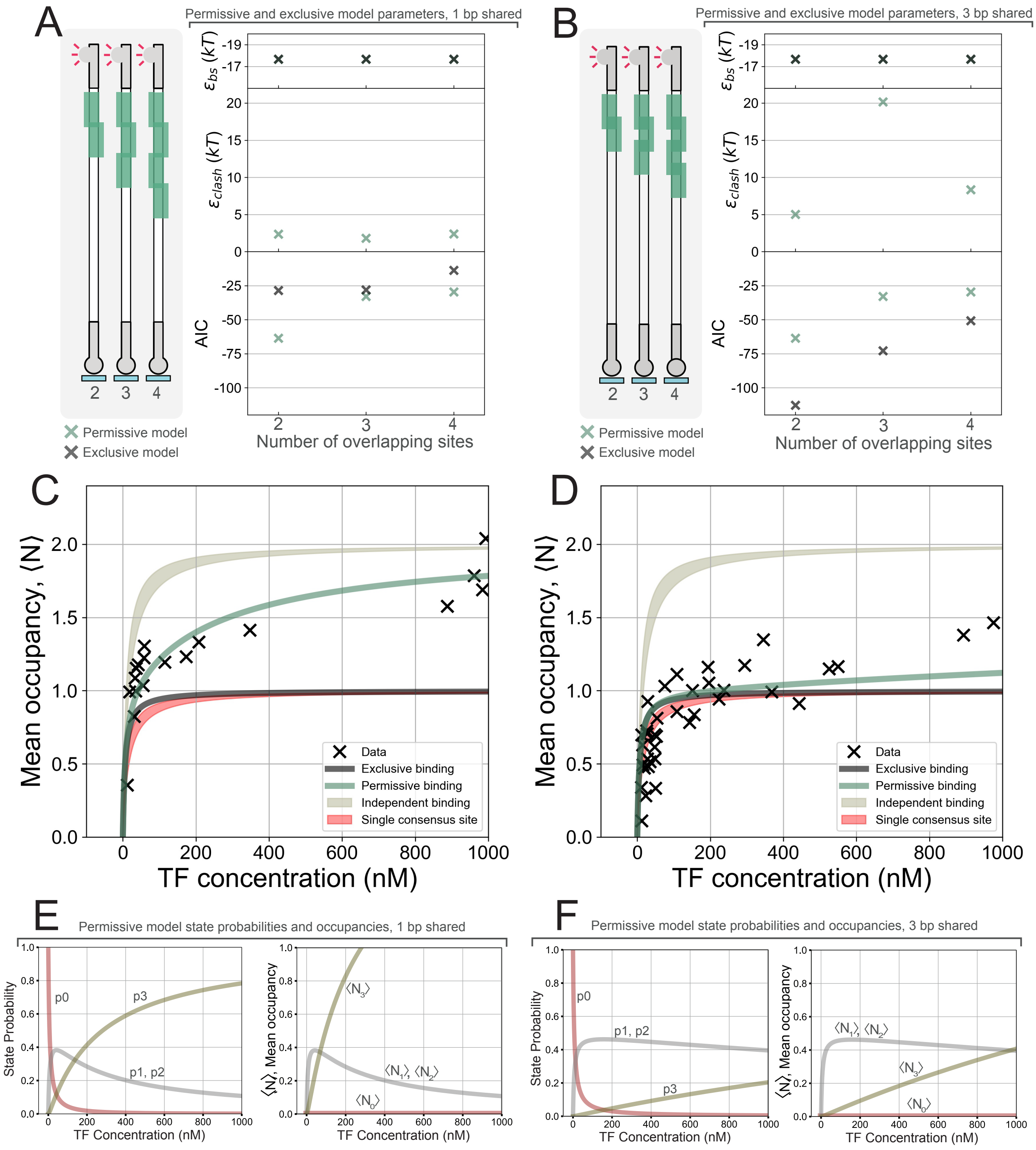

Figure S5: Modeling of binding to high-density clusters.

Caption on the next page.

Figure S5: (**A**) DNA targets for high-density clusters with 1 shared basepair betweeen neighboring consensus binding sites (left). Corresponding parameters and Akaike information criteria (AIC) for exclusive and permissive models of binding (right). Energies are reported in units of $kT$ (product of Boltzmann constant and temperature). (**B**) DNA targets (left) and model parameters (right) for high-density clusters with 3 shared basepairs between neighboring consensus binding sites. **(C)** Characterization data for the DNA target in (**A**) with two consensus binding sites sharing 1 basepair. Exclusive and permissive model predictions in grey and green respectively. 95% confidence intervals for the characterized single consensus binding site (red), and a model of independent binding to two consensus binding sites (beige). (**D**) Corresponding data and models for the DNA target in **(B)** with two consensus binding sites sharing 3 basepairs. (**E, F**) State probabilities and occupanices for the permissive model, for the 2 consensus site DNA targets with 1 shared basepair **(E)** and with 3 shared basepairs **(F)**.

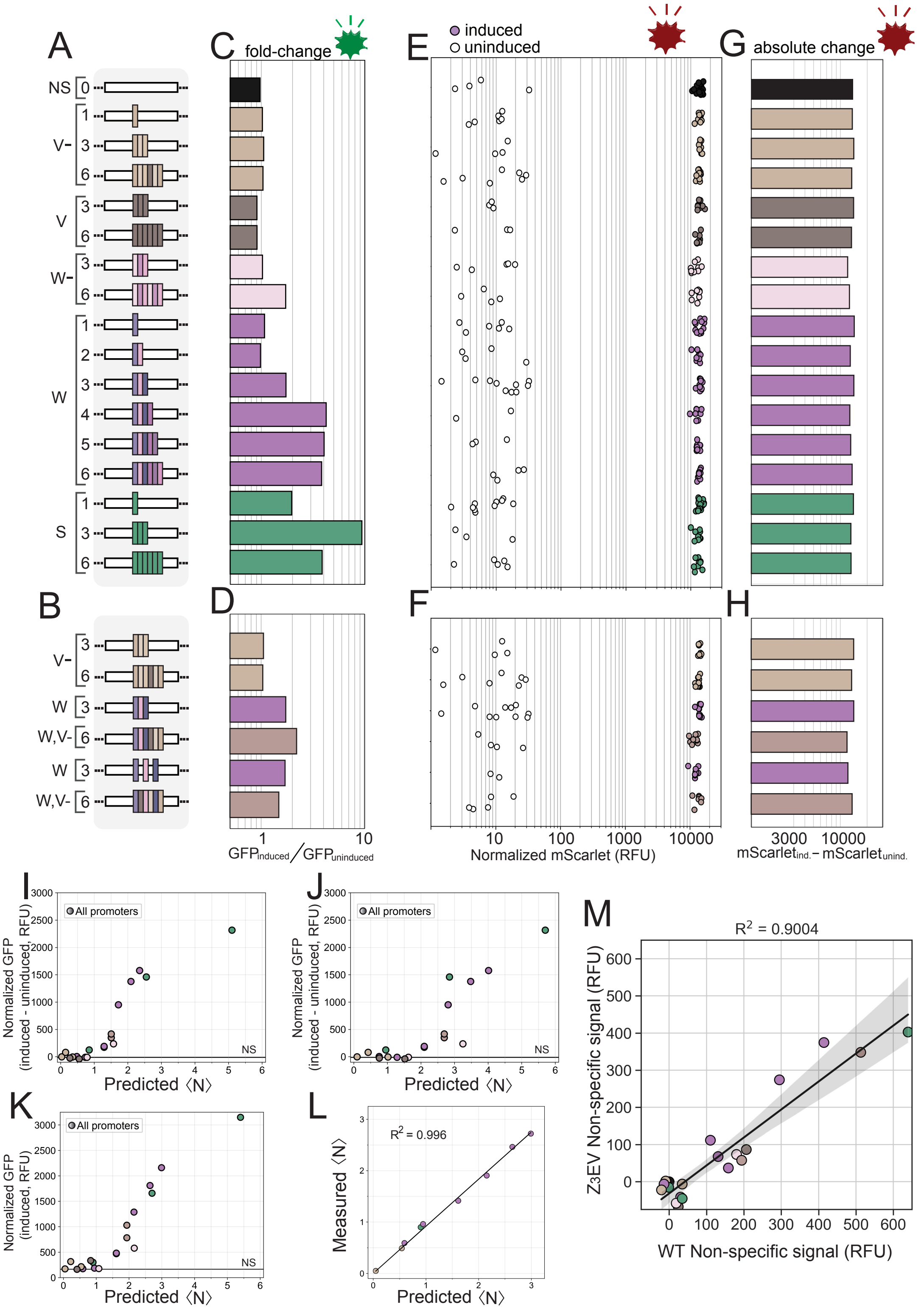

Figure S6: Alternate analyses and control data for the Zif268 *in vivo* results in Figures 5 and 6. Caption on the next page.

Figure S6: **(A, B)** Legends for the characterized minCYC1 promoter libraries. **(C, D)** Alternate data analysis to Figure 5G and 5H, quantifying the Zif268-specific impact on gene expression as GFP mean fold-change between induced and uninduced strains. **(E, F)** Induced and uninduced mScarlet expression levels driven by the control promoter. **(G, H)** Absolute difference between induced and uninduced mScarlet expression levels. **(I, J)** Alternate data analysis to Figure 6, varying the transcription factor concentration from 73 nM (85% consensus saturation) **(I)** to 245 nM (95% consensus saturation) **(J)**. **(K)** Changing the measure of specific gene expression from the difference between mean induced and mean uninduced GFP signal, to mean induced GFP signal. **(L)** Relationship between predicted mean occupancy and measured mean occupancy, for the *in vitro*-characterized DNA targets that were used in the Zif268 *in vivo* study. Predictions were based on an independent statistical thermodynamic binding model similar to Equation 24, but with a variable number of binding sites, parameterized by data collected through characterization of single-site DNA targets, as explained in equation 6.4. $R^2$ of 0.996. **(M)** Relationship between the non-specific signal for the $Z_3$EV strains containing the $Z_3$EV transcription factor, and the WT (BY4741) strain containing no transcription factors with a Zif268 DNA binding domain. In each case the non-specific signal was quantified as the absolute mean difference between a given promoter's GFP expression and the GFP expression of the nonspecific (NS) promoter containing no binding sites (similar to Figure 5I and 5J), both in the respective strain's background. $R^2$ of 0.9, suggesting that the expression in uninduced strains is due to an endogenous transcription factor.

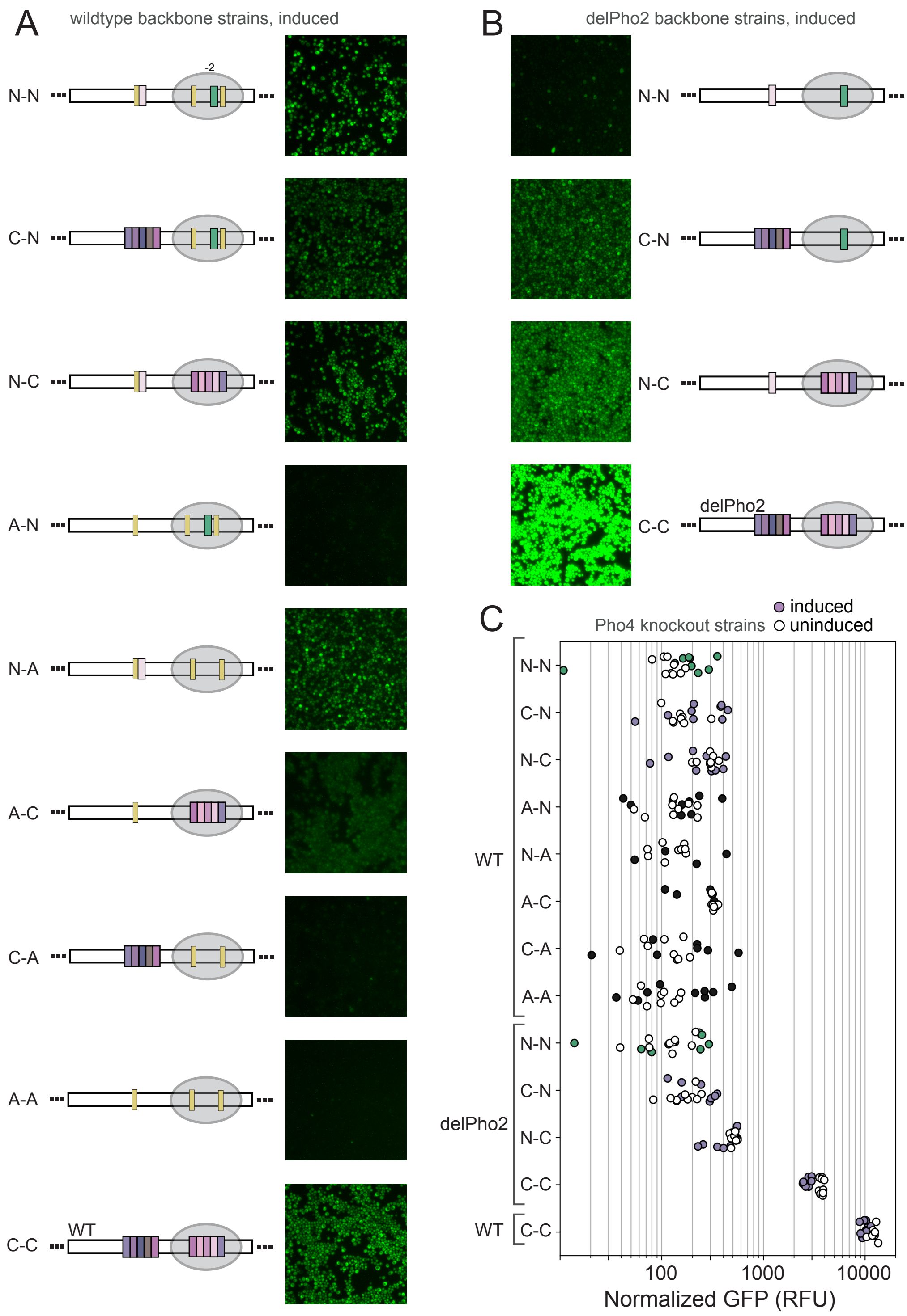

Figure S7: Fluorescence microscope images for the BY4741 strain library, and induced and uninduced platereader measurements for the Pho4 knockout strains.

Caption on the next page.

Figure S7: **(A)** Fluorescence microscope images of the BY4741 strains with wildtype PHO5 promoter backbones induced by phosphate starvation. **(B)** Fluorescence microscope images of the BY4741 strains with delPho2 (all Pho2 sites ablated) PHO5 promoter backbones induced by phosphate starvation. **(C)** Platereader measurements following induction or in the absence of induction, for Pho4 knockout strains, enlarged from Figure 7L. Platereader data was processed similarly to the other platereader measurements in Figure 7.

### Part II

### Materials and methods

#### 1 Purified protein and iMITOMI DNA target production

##### 1.1 DNA target PCRs

Initial materials for producing iMITOMI DNA target libraries included tagged primers (5' Cy5-tagged: CTG/iCy5/TCGGCCGCTAACA, and 3' biotin-tagged: /5Biosg/GTCATACCGCCGGA) ordered from IDT. Furthermore, DNA targets of interest were ordered as single-stranded 90-basepair long primers from IDT, which had 5' and 3' ends complementary to the tagged primers. For each library member, two 50uL PCR reactions were prepared according to the following recipe:

| Reagent | Initial conc. | Volume to add (per 50 $\mu$L rxn) |
|---|---|---|
| Biotin-tagged primer | 100 $\mu$M | 0.25 $\mu$L |
| Cy5-tagged primer | 100 $\mu$M | 0.25 $\mu$L |
| 90bp ssDNA target primer | 500 nM | 0.25 $\mu$L |
| DreamTaq Green PCR Master Mix | 2X | 25 $\mu$L |
| Invitrogen UltraPure Distilled Water | | 24.25 $\mu$L |

Table S1: iMITOMI targets' PCR recipe

| Step | Temperature | Duration |
|---|---|---|
| 1. | 95°C | 1 minute |
| 2. | 95 °C | 30 seconds |
| 3. | 50°C | 30 seconds |
| 4. | 72°C | 1 minute |
| ***Steps 2 to 4 are cycled a total of 30 times*** | | |
| 5. | 72°C | 10 minutes |
| 6. | 4°C | ∞ |

Table S2: iMITOMI targets' PCR cycling conditions

Two 50 $\mu$L PCR products were mixed with 500 $\mu$L of DNA binding buffer, combined into one purification column (Zymo DNA Clean & Concentrator 50), and PCR purified according to kit instructions. Samples were eluted into 80 $\mu$L of Invitrogen UltraPure Distilled Water. DNA targets were run on an agarose gel for characterization, and only products with a single band of expected size were advanced to spotting.

#### 1.2 Spotting plate preparation

75 $\mu$L of each DNA target was introduced into independent wells of a 384 well plate. Plates were measured for fluorescence with an excitation wavelength of 570±9 nM and an emission wavelength of 593±9 nM, using a Biotek Synergy Mx Multi-mode reader. Based on these measurements, new fluorescence-equalized wells were prepared by diluting an appropriate amount of DNA with water to reach 72 $\mu$L (no water was added for the well with the lowest fluorescence intensity, the others were equalized to this fluorescence intensity ± 5%, Supp. Figure S1C. 24 $\mu$L of 2% Bovine Serum Albumin (in $H_2O$) was added to each well. The plate was sealed with aluminum and kept at -20°C until time of use.

#### 1.3 His-tag protein purification

For each transcription factor, bacteria containing the corresponding plasmid were overnight cultured with ampicilin (Amp), and innoculated into two 1 L baffled flasks each containing 500 mL LB and

Amp (100 ug/ml). Cultures were grown at 37°C at 260 rpm for 2 hours, induced with IPTG (1 mM) and grown for an additional 3 hours. Cells were centrifuged at 3220 g at 4°C for 10 min in 50 mL falcon flasks. Cell pellets were resuspended in buffer A (sequential transfers, using total volume of 7.5 mL), and sonicated (Vibra cell 75186 sonicator, probe tip diameter: 6 mm) with 20 s ON 20 s OFF for four cycles using 70% amplitude. Lysate was distributed to 2 mL tubes and centrifuged for 20 minutes at 4°C at 20,000 g. Supernatant was removed and dispensed into a regenerated Ni Sulfate column. The column was washed with 25 mL buffer A, then protein was eluted in 5 mL buffer B. The sample was dialysed overnight in 1 L of HT buffer at 4°C with magnetic stirring, followed by 3 hours of dialysis with HT stock buffer under similar conditions. The sample was run on an SDS-PAGE gel in S2 for characterization.

#### 1.4 Purification buffer recipes

| **Component** | **Final conc.** |
|---|---|
| **Buffer A** | |
| NH4Cl | 1 M |
| HEPES | 50 mM |
| MgCl | 10 mM |
| B-ME (fresh) | 7 mM |
| **Buffer B** | |
| Imidazole | 0.5 M |
| HEPES | 50 mM |
| MgCl | 10 mM |
| KCl | 100 mM |
| B-ME (fresh) | 7 mM |
| **HT buffer** | |
| HEPES | 50 mM |
| MgCl | 10 mM |
| KCl | 100 mM |
| B-ME (fresh) | 7 mM |

| HT stock buffer | |
|---|---|
| HEPES | 50 mM |
| MgCl | 10 mM |
| KCl | 100 mM |
| Glycerol | 60% |
| B-ME (fresh) | 7 mM |

Table S3: Purification buffer recipes

#### 2 Control and flow wafer fabrication

Mold-fabrication using silicon wafers was performed through a photolithography process in the CMi cleanrooms at EPFL, as described previously (Laohakunakorn et al. (2019); Maerkl and Quake (2007b)). In brief, for the control layer, a silicon wafer was primed in a Tepla 300, and SU-8 photoresist (GM 1070) was spin coated onto the wafer using a Sawatec spin coater to reach a height of 30 µm. After a soft bake the wafer was exposed (365 nm illumination, 20 mW/cm$^2$ light intensity) using a chrome mask for 10 s on a MABA6 mask aligner. The wafer was post exposure baked and then developed using propylene glycol methyl ether acetate (PGMEA), followed by a hard bake. For the flow layer the silicon wafer was treated with hexamethyldisilizane (HMDS) vapor using a YesIII oven. AZ 9260 photoresist was spin coated onto the wafer using an EVG150 modular cluster tool to reach a height of around 14 µm. After the wafer was baked it was left for a one hour relaxation period, followed by UV exposure using an MABA6 mask aligner. The total dose was 660 mJ/cm split into two exposures of 18s with a 10s wait period in between (20 mW/cm$^2$ light intensity). The wafer was developed again using the EVG150 (AZ 400K developer), and then baked at 160 °C for two hours.

Designs of microfluidic devices are available on our group's website: http://lbnc.epfl.ch.

### 3 PDMS chip fabrication

Wafers were first treated with chlorotrimethylsilane (TMCS) vapor to facilitate removal of cured polydimethylsiloxane (PDMS) from the wafers. PDMS curing agent and elastomer (amounts of 1 g and 20 g for the flow layer, 10 g and 50 g for the control layer, respectively) were mixed (2000 rpm for 1 min) and defoamed (2200 rpm for 2 min) using a Thinky ARE-250 centrifugal mixer. PDMS from the control layer was poured on the control wafer and left to degas in a vacuum chamber, followed by baking for 20 minutes. For the flow layer, 4 mL of PDMS was spin-coated onto the flow-wafer using a SCS G3P-8 spin coater (1800 rpm for 35s), followed by baking for 20 minutes. Control inlets were punched using a Schmidt Press manual hole puncher and 21-gauge (OD 0.04") pins (Technical Innovations, Inc.), and the control PDMS blocks were aligned to the flow layer under a stereoscope using coaxial illumination. Aligned chip layers were bonded at 80°C for 90 minutes. Next, PDMS chips were cut off the flow layer, and flow inlets and outlets were punched using a 900 mm pin.

### 4 Epoxy slide coating and microarraying

VWR slides with cut edges and plain ends (631-1550) were cleaned in a solution of 600 mL miliQ water, 120 mL 25% Ammonia solution, and 150 mL $H_20_2$ (on a hotplate at 80°C). Slides were then washed with miliQ water and dried. Next, slides were left to incubate for 20 minutes in a bath of 891 mL toluene and 5 mL of (3-Glycidyloxypropyl)trimethoxysilane (GPS, Sigmaaldrich Cat. 440167). Slides were rinsed with toluene and dried, followed by baking at 120°C for 120 minutes.

Epoxy-coated glass slides were spotted with the purified 90 bp DNA targets using a Genetix Qarray$^2$ robot with an MP2.5 pin (Arrayit). Spotting chambers were aligned to DNA spots on the glass slide under a microscope, and the device was baked for 8 hours at 80°C.

### 5 iMITOMI experiments

Control lines were filled with tap water and pressurized at 172 kPa. The the neck valve was pressurized to prevent premature target resolubilization. Flow lines were pressurized at 48 kPa. Surface chemistry was conducted in the remainder of the chip (Maerkl and Quake (2007a)), by sequentially patterning the surface using 15 minute flow steps with biotin-BSA, then neutravidin, followed by actuation of the button valve and a second biotin-BSA flow. PBS washes were conducted between each flow step for 5 minutes. This resulted in available neutravidin binding sites under the button valve in the detection chamber, whereas the remainder of neutravidin binding sites in the chip's flow channels were passivated by biotin-BSA. DNA targets in spotting chambers were resolubilized by pressurizing the PBS flow line, closing the outlet and opening the neck valve to force air out of the PDMS and PBS into the spotting chambers. After a PBS wash with the neck valve closed to prevent any contamination, the sandwich valves were pressurized to isolate chambers from one another, and the neck and button valves were opened to allow the biotinylated Cy5-tagged DNA targets to diffuse to and bind under the button valves in the detection chambers for 90 minutes. The button valve was then closed, and the chip was washed with PBS. The chip surface was blocked using Promega wheat-germ extract to reduce non-specific binding of concentrated fluorescent protein solutions. The chip was multiplexed into 8 sections using the multiplexing valves. Different concentrations of purified mScarlet-tagged transcription factor were flowed into different sections of the chip, followed by incubation for 60 minutes with the button valves open to allow binding to reach equilibrium. The chip was scanned to quantify the free protein concentration at equilibrium (compared against a background scan after the subsequent wash step). The button valve was closed to isolate the bound protein at equilibrium, and the chip was washed, and scanned again to quantify the bound protein and amount of DNA present (compared against a background scan before fluorescent protein or fluorescent DNA was introduced into the detection chamber, respectively).

### 6 iMITOMI data analysis and modeling

#### 6.1 Image analysis and data processing

Images from a given scan were stitched together using ImageJ and ROIs were processed using GenePix, which facilitated automatic and manual feature alignment, as well as conversion of ROI pixel intensities to mean signal intensities.

#### 6.2 Quantification of free and bound protein signals

A normalized bound signal intensity was quantified as:

$$S_{bound} = \frac{TF_{bound} - TF_{background}}{DNA_{immobilized} - DNA_{background}} \tag{1}$$

$S_{bound}$ corresponds to the bound transcription factor at equilibrium (background corrected) normalized by the amount of DNA at equilibrium (background corrected). The values used in this calculation were mean signal intensities from an ROI under the button valve, at the appropriate assay stage discussed in section 5.

A signal $S_{free}$ corresponding to free protein at equilibrium was quantified using an ROI next to the button valve as:

$$S_{free} = TF_{free} - TF_{background} \tag{2}$$

This signal was calibrated against the calibration curves from Figure S2 to obtain absolute TF concentrations, [TF].

#### 6.3 Saturation binding curve parameter estimation

According to mass action kinetics, at equilibrium a 1:1 transcription factor (TF) to binding site (BS) binding reaction follows the relationship

$$[\mathrm{TF}] + [\mathrm{BS}] \underset{\mathrm{k}_{off}}{\overset{\mathrm{k}_{on}}{\rightleftharpoons}} [\mathrm{TF{:}BS}]$$

$$k_{on} \cdot [TF][BS] = k_{off} \cdot [\mathrm{TF{:}BS}]$$

$$K_D = \frac{k_{off}}{k_{on}} = \frac{[TF][BS]}{[\mathrm{TF{:}BS}]}$$

$$[\mathrm{TF{:}BS}] = \frac{[TF][BS]}{[K_D]} \tag{3}$$

Furthermore, the proportion of binding sites bound by transcription factor molecules, $P_{bound}$, can be defined as follows

$$P_{bound} = \frac{[\mathrm{TF{:}BS}]}{[\mathrm{TF{:}BS}] + [BS]} \tag{4}$$

Through substituting Equation 3 into Equation 4 and rearranging, it follows that

$$P_{bound} = \frac{[TF]}{K_D + [TF]} \tag{5}$$

This derives a relationship to model the proportion of binding sites bound by transcription factor, based on the concentration of free transcription factor and the binding reaction's dissociation constant ($K_D$, representing the affinity).

The same functional form as in Equation 5 can be used similarly to describe a binding reaction to DNA targets containing multiple binding sites, if all binding site affinities are assumed equivalent, and binding is assumed to be independent. To see this, it is useful to consider that [BS] was used in the above analysis instead of [DNA], and the analysis is not sensitive to if some constant number of [BS] are distributed on fewer molecules of DNA. The $B_{max}$ parameter represents the fluorescence intensity at transcription factor concentrations for a given DNA target, and using it to scale $P_{bound}$ allows for a rescaling from proportion of binding sites bound to $S_{bound}$, that is

$$S_{bound} = P_{bound} \cdot B_{max} \tag{6}$$

Therefore, experimental data characterizing saturation binding to a given DNA target can be fit according to the two parameter ($K_D$, $B_{max}$) model in equation 7

$$S_{bound} = \frac{B_{max} \cdot [TF]}{K_D + [TF]} \tag{7}$$

where [TF] is in absolute concentration units (nM) to obtain a $K_D$ estimate in nM. As discussed, $B_{max}$ represents the fluorescence level at saturating transcription factor concentrations.

In order to model cooperative binding, a common phenomenological modification to Equation 7 is the incorporation of the Hill coefficient ($n_H$), even for 1:1 binding reactions, which produces the Hill model:

$$S_{bound} = \frac{B_{max} \cdot [TF]^{n_H}}{K_D + [TF]^{n_H}} \tag{8}$$

For a given DNA target, data were combined across two to six experimental days and an initial fit to Equation 7 was obtained through non-linear least squares minimization with the Lmfit package in Python (Newville et al. (2014)). To explore the parameter space around this fit initial parameter set and obtain confidence intervals, we used Markov Chain Monte Carlo to conduct sampling of the posterior probability distribution given the experimental data and assuming a uniform prior (Figure S4B and C).

To obtain mean occupancies $\langle N \rangle$ , $S_{bound}$ (units RFUs) was normalized (calibrated) according to the $B_{max}$ increase from adding additional binding sites (or equivalently, the RFUs per 1 TF molecule bound to DNA on average), as described in the results section of the main text. This converts equation 7 to the more interpretable form in equation 9

$$\langle N \rangle = \frac{n_{sites} \cdot [TF]}{K_D + [TF]} \tag{9}$$

where the mean occupancy $\langle N \rangle$ represents the average number of transcription factor molecules bound per DNA molecule, and $n_{sites}$ represents the number of defined binding sites on the DNA target. Upon normalization, $B_{max}$ on the right hand side is converted to $n_{sites}$ since the $B_{max}$ of a given target can be thought of as the $B_{max}$ from each binding site, multiplied by the number of binding sites.

#### 6.4 Statistical mechanical modeling

Statistical mechanical models of binding were built starting from the useful framework developed by Philips et. al. and presented in the Physical Biology of the Cell textbook (Phillips et al. (2012)). We recommend referring to the textbook for a detailed discussion of the modeling framework. Essentially, microstates of the binding system were grouped into states, where each state represents all of the possible microstates of the system where a particular set of DNA binding sites are bound by transcription factor molecules (e.g. the unbound state represents all microstates of the system where the DNA is not bound by any transcription factor molecule). Here a microstate signifies a particular distinguishable arrangement of indistinguishable transcription factor molecules among positions of a spatially discretized environment (e.g. the unbound state is composed of many microstates, which differ from one another based on which positions in the lattice-modeled solution are occupied). Each state can be attributed a total energy, by accounting for the energies from all particles in one of its microstates. This total energy does not differ between the different microstates of a given state. Furthermore, each state is attributed a multiplicity, by counting the number of different microstates that can give rise to the state.

In order to calculate the probability of any state, first recognize that

$$p_i \propto e^{-E_i/(kT)}$$

$$\beta = \frac{1}{kT}$$

$$p_i \propto e^{-\beta E_i} \tag{10}$$

where $p_i$ is the probability of microstate $i$, $E_i$ is the total energy from all particles in microstate $i$, and the constant $kT$ is a product of the Boltzmann constant $k$ and the absolute temperature $T$ and usually written as $\beta$. The term $e^{-\beta E_i}$ is the Boltzmann factor. Equation 11 relates the ratio of probabilities for two different microstates to the ratio of their Boltzmann factor's, which depends on the difference between their total energies. This captures the typical intuition that lower-energy configurations are more favorable.

$$\frac{p_i}{p_j} = e^{\beta(E_j - E_i)} \tag{11}$$

To determine the probability for a given microstate (Equation 12), its Boltzmann factor must be normalized by the sum of the Boltzmann factors from every possible microstate (the partition function, Z, Equation 13), such that the microstate probabilities sum to 1.

$$p_i = \frac{e^{-\beta E_i}}{Z} \tag{12}$$

$$Z = \Sigma_i \ e^{-\beta E_i} \tag{13}$$

In order to calculate the probability of any given state (group of microstates), it follows that the state's multiplicity can be multiplied by the state's Boltzmann factor. This produces what is effectively the sum over Boltzmann factors from all microstates belonging to the state, a term referred to as the state's weighted multiplicity (Phillips et al. (2012)), which accordingly when normalized by the partition function produces the state's probability.

For illustration, it is useful to first consider how to count the number of microstates (how to calculate the multiplicity) for the unbound state

$$\frac{\Omega!}{L!(\Omega - L)!} \tag{14}$$

Where $\Omega$ represents the number of discrete positions where a transcription factor could be located in solution, and $L$ represents the number of transcription factor molecules. Transcription factor molecules can be placed in solution in $\frac{\Omega!}{(\Omega-L)!}$ different configurations, but they can be rearranged indistinguishably $L!$ ways.

Furthermore, the following related approximation will enable us to simplify our analysis

$$\frac{x!}{y!(x-y)!} \approx \frac{x^y}{y!}, \quad \text{if } x \gg y \tag{15}$$

More generally, state multiplicites evaluate to

$$\frac{\Omega!}{(L-n)!\ (\Omega-(L-n))!} \tag{16}$$

Where $n$ represents the number of TF molecules bound to the DNA molecule

Which according to Approximation 15 simplifies to

$$\frac{\Omega^{L-n}}{(L-n)!} \tag{17}$$

Since we assume that in our binding system $\Omega \gg L$

Table S4 and Equations 18 to 25 demonstrate how to model a simple system of transcription factor molecules binding to DNA having a single binding site, in order to illustrate how the model in Equation 5 that we derived from mass action kinetics can be equivalently derived through statistical mechanics. This connection ultimately allowed us to coherently integrate both mass action kinetic and statistical mechanical modeling in our study.

| State | Energy, $E_i$ | Multiplicity | Relative weighted multiplicity |
|---|---|---|---|
| $\underline{\ \ }$ | $L \cdot \varepsilon_{sol}$ | $\frac{\Omega!}{L!(\Omega-L)!} \approx \frac{\Omega^L}{L!}$ | 1 |
| $\underline{O}$ | $\varepsilon_{bs} + (L-1) \cdot \varepsilon_{sol}$ | $\frac{\Omega!}{(L-1)!(\Omega-(L-1))!} \approx \frac{\Omega^{L-1}}{(L-1)!}$ | $\frac{[TF]}{[TF]_o} \cdot e^{-\beta\Delta\varepsilon_{bs}}$ |

Table S4: 1:1 binding through the lens of statistical mechanics

The total energy, $E_i$, of the unbound state ( $\underline{\ \ }$ ) is signified by the sum of an $\varepsilon_{sol}$ energy term from each of the $L$ particles in solution. For the bound state ( $\underline{O}$ ), one particle less will be at an energy of $\varepsilon_{sol}$, and instead will have the energy $\varepsilon_{bs}$, representing its specific interaction to a single binding site on DNA. Multiplicities are weighted according to their Boltzmann factors as follows, for the unbound state the weighted multiplicity is

$$\frac{\Omega^L}{(L)!} \cdot e^{-\beta L \varepsilon_{sol}} \tag{18}$$

And for the bound state, the weighted multiplicity is

$$\frac{\Omega^{L-1}}{(L-1)!} \cdot e^{-\beta((L-1)\varepsilon_{sol}+\varepsilon_{bs})} \tag{19}$$

Furthermore, following the convention in (Phillips et al. (2012)), we normalize the weighted multiplicities and partition function by the weighted multiplicity of the unbound state, to represent these terms as relative weighted multiplicities (Table S4, Equation 20). For the bound state the relative weighted multiplicity simplifies to

$$\frac{\frac{\Omega^{L-1}}{(L-1)!} \cdot e^{-\beta((L-1)\varepsilon_{sol}+\varepsilon_{bs})}}{\frac{\Omega^L}{(L)!} \cdot e^{-\beta L \varepsilon_{sol}}}$$

$$= \frac{L}{\Omega} \cdot e^{-\beta(\varepsilon_{bs}-\varepsilon_{sol})} \tag{20}$$

Furthermore, as illustrated in Phillips et al. (2012),

$$[TF] = \frac{L}{\Omega \cdot V_{box}}$$

$$[TF]_o = \frac{1}{V_{box}}$$

$$\frac{[TF]}{[TF]_o} = \frac{L}{\Omega} \tag{21}$$

where [TF] is the concentration of transcription factor, and the constant $[\mathrm{TF}]_o$ is a reference concentration that corresponds to if each position in solution was occupied by a transcription factor molecule (Phillips et al. (2012)). Furthermore, $\mathrm{V}_{box}$ is the volume of one discrete lattice position in solution. This allows the bound state's relative multiplicity to be expressed as

$$\frac{[TF]}{[TF]_o} \cdot e^{-\beta \Delta\varepsilon_{bs}}, \quad \text{where } \Delta\varepsilon_{bs} = \varepsilon_{bs} - \varepsilon_{sol} \tag{22}$$

Then, the probability of any state can be determined by normalizing by the partition function (expressed relative to the unbound state) Z, which is the sum of all relative weighted multiplicities

$$Z = 1 + \frac{[TF]}{[TF]_o} \cdot e^{-\beta \Delta \varepsilon_{bs}} \tag{23}$$

Therefore, the probability of the bound state is

$$p_{bound} = \frac{\frac{[TF]}{[TF]_o} \cdot e^{-\beta \Delta \varepsilon_{bs}}}{1 + \frac{[TF]}{[TF]_o} \cdot e^{-\beta \Delta \varepsilon_{bs}}} \tag{24}$$

We used the single-site binding model in Equation 24, which is equivalent to the expression for mean occupancy for single-site targets, to obtain energy parameters for particular binding sites, by fitting the corresponding single-site DNA targets' binding data (normalized by the increase in $B_{max}$ with additional sites) to the model (using non-linear least-squares minimization, followed by MCMC, as before). These binding site energies were used to parameterize higher-order statistical mechanical models of binding to DNA targets containing multiple binding sites (e.g. for modeling exclusive and permissive binding to high-density clusters).

Equation 24 is the statistical mechanical equivalent to Equation 5, considering that

$$K_D = [TF]_o \cdot e^{\beta \Delta \varepsilon_{bs}} \tag{25}$$

As discussed, this connection permitted us to move between statistical mechanical and mass action modeling frameworks in our study.

#### 6.5 Exclusive binding model to describe binding to high-density clusters

In exclusive binding models, transcription factors were not permitted to be bound simultaneously to overlapping binding sites (sites sharing common basepairs), as illustrated in the main manuscript Figure 4D and Table S5. For example, for a cluster with two binding sites, only three states (unbound, left site bound, or right site bound) were considered in the exclusive model. The states, energies, and multiplicites for an exclusive model of binding to a DNA target containing two binding sites are shown in Table S5.

| State | Energy, $E_i$ | Multiplicity | Relative weighted multiplicity |
|---|---|---|---|
| — | $L \cdot \varepsilon_{sol}$ | $\frac{\Omega!}{L!(\Omega-L)!} \approx \frac{\Omega^L}{L!}$ | 1 |
| O_ | $\varepsilon_{bs1} + (L-1) \cdot \varepsilon_{sol}$ | $\frac{\Omega!}{(L-1)!(\Omega-(L-1))!} \approx \frac{\Omega^{L-1}}{(L-1)!}$ | $\frac{[TF]}{[TF]_o} \cdot e^{-\beta \Delta\varepsilon_{bs1}}$ |
| _O | $\varepsilon_{bs2} + (L-1) \cdot \varepsilon_{sol}$ | $\frac{\Omega!}{(L-1)!(\Omega-(L-1))!} \approx \frac{\Omega^{L-1}}{(L-1)!}$ | $\frac{[TF]}{[TF]_o} \cdot e^{-\beta \Delta\varepsilon_{bs2}}$ |

Table S5: Exclusive binding model for a high-density cluster with two overlapping sites

In order to calculate the mean occupancy of transcription factors on DNA, we can calculate a probability-weighted average of the number of transcription factors bound in each state. Therefore, for the example of exclusive binding to two overlapping sites (states presented in Table S5)

$$\langle N \rangle = 0 \times \frac{1}{Z} + 1 \times \frac{\frac{[TF]}{[TF]_o} \cdot e^{-\beta\Delta\varepsilon_{bs1}}}{Z} + 1 \times \frac{\frac{[TF]}{[TF]_o} \cdot e^{-\beta\Delta\varepsilon_{bs2}}}{Z}, \quad \text{where} \tag{26}$$

$$Z = 1 + \frac{[TF]}{[TF]_o} \cdot e^{-\beta\Delta\varepsilon_{bs1}} + \frac{[TF]}{[TF]_o} \cdot e^{-\beta\Delta\varepsilon_{bs2}} \tag{27}$$

#### 6.6 Permissive binding model to describe binding to high-density clusters

In permissive binding models, transcription factors were permitted to be bound simultaneously to overlapping binding sites, as illustrated in the main manuscript Figure 4D and Table S6. For example, for a cluster with two overlapping binding sites, four states were permitted (unbound, left site bound, right site bound, or both sites bound).

| State | Energy, $E_i$ | Multiplicity | Relative weighted multiplicity |
|---|---|---|---|
| — | $L \cdot \varepsilon_{sol}$ | $\frac{\Omega!}{L!(\Omega-L)!} \approx \frac{\Omega^L}{L!}$ | 1 |
| O_ | $\varepsilon_{bs1} + (L-1) \cdot \varepsilon_{sol}$ | $\frac{\Omega!}{(L-1)!(\Omega-(L-1))!} \approx \frac{\Omega^{L-1}}{(L-1)!}$ | $\frac{[TF]}{[TF]_o} \cdot e^{-\beta \Delta\varepsilon_{bs1}}$ |
| _O | $\varepsilon_{bs2} + (L-1) \cdot \varepsilon_{sol}$ | $\frac{\Omega!}{(L-1)!(\Omega-(L-1))!} \approx \frac{\Omega^{L-1}}{(L-1)!}$ | $\frac{[TF]}{[TF]_o} \cdot e^{-\beta \Delta\varepsilon_{bs2}}$ |
| OO | $\varepsilon_{bs1} + \varepsilon_{bs2} + \varepsilon_{clash} + (L-2) \cdot \varepsilon_{sol}$ | $\frac{\Omega!}{(L-2)!(\Omega-(L-2))!} \approx \frac{\Omega^{L-2}}{(L-2)!}$ | $(\frac{[TF]}{[TF]_o})^2 \cdot e^{-\beta(\Delta\varepsilon_{bs1}+\Delta\varepsilon_{bs2}+\varepsilon_{clash})}$ |

Table S6: Permissive binding model for a high-density cluster with two overlapping sites

For binding to larger clusters where two or more transcription factors can be bound at once (i.e. the state with both sites bound in the permissive model in Table S6), this must be taken into account in these states' multiplicities. For illustration, consider the multiplicity of a state with two transcription factors bound at once. According to Equation 16 and Equation 17.

$$\frac{\Omega!}{(L-2)!\ (\Omega-(L-2))!} = \frac{\Omega^{L-2}}{(L-2)!} \tag{28}$$

Which when normalized by the multiplicity of the unbound state results in the relative multiplicity

$$\frac{\frac{\Omega^{L-2}}{(L-2)!}}{\frac{\Omega^{L}}{L!}} = \frac{L(L-1)}{\Omega^2} \tag{29}$$

The term on the RHS of Equation 29 can be re-written as

$$\frac{\frac{L!}{(L-2)!}}{\Omega^2} \tag{30}$$

Which according to Approximation 15

$$= \frac{L^2}{\Omega^2} \quad if\ \ L \gg 2 > 0 \tag{31}$$

More generally, a state with $n$ transcription factors bound at once will have the relative weighted multiplicity

$$\left(\frac{L}{\Omega}\right)^n, \quad \text{if } L \gg n > 0$$

$$= \left(\frac{[TF]}{[TF]_o}\right)^n \tag{32}$$

Permissive model states possess an additional energy parameter $\varepsilon_{clash}$ (TableS6, bottom) for each set of shared basepairs (at the junction of two overlapping sites) where two transcription factors are bound simultaneously. For instance, for a cluster with three binding sites where the first site overlaps the second, and the second overlaps the third, the state with three transcription factors bound simultaneously would possess $2 \cdot \varepsilon_{clash}$in its energy term. $\varepsilon_{clash}$ can be considered as composed of both an energy associated with steric interference between transcription factor molecules, and some loss of binding to the shared basepairs, lumped into a single parameter.

#### 6.7 Model selection

To compare models of binding to high-density clusters, the Akaike Information Criterion (AIC) and Bayesian Information Criterion (BIC) were computed using the Lmfit package (Newville et al. (2014)). The AIC and BIC can be computed as:

$$AIC = N \cdot ln(\frac{\chi^2}{N}) + 2N_{vars} \tag{33}$$

$$BIC = N \cdot ln(\frac{\chi^2}{N}) + ln(N) \cdot N_{vars} \tag{34}$$

$$\tag{35}$$

Where $\chi^2$ is the chi-square, $N$ is the number of data points, and $N_{vars}$ is the number of floating parameters. For clusters where 1 basepair was shared between neighboring binding sites, the lower AIC and BIC suggest that the permissive model better represents binding than the exclusive model (Figure S5). On the other hand, as the number of shared basepairs increased to 33% of the motif (3bp), binding became largely exclusive, as reflected by the lower AIC and BIC for the exclusive model.

### 7 *In vivo* data analysis and modeling

#### 7.1 Modeling independent binding to clusters using parameters derived from single site DNA targets

Our results identified that binding sites with a positive gap distance in lower-density clusters exhibit largely independent binding. Therefore to model binding to these clusters, accounting for the difference in affinities between different sites, we used an independent binding model parameterized by energies derived from characterization of the individual binding sites on single site DNA targets. We used a statistical mechanical model that accounts for the different states of the system, although a similar result could be obtained by summing the occupancy contribution from each individual binding site in isolation.

To predict the mean occupancy of Zif268 clusters *in vivo*, we simulated a concentration range from 85% to 95% saturation of the consensus binding site (73 nM to 245 nM), confirming our results under a wide range of concentrations expected for the $Z_3$EV transcription factor. To check the accuracy of our model and premise of binding independence, for clusters that we characterized fully *in vitro* we compared our model predictions to measured occupancies (Figure S6M), which showed near perfect agreement and an $R^2$ value of 0.996.

To quantify induced and uninduced levels of gene expression from platereader timeseries measurements, the signal in the appropriate fluorescence or OD600 channel from blank wells containing media alone was subtracted. Then the fluorescence timeseries measurements were normalized by their corresponding OD600 values. To represent induced and uninduced levels for each replicate, timeseries measurements were averaged across a time window before the strains reached stationary phase, and all data was plotted in strip plots using Seaborn and Matplotlib in Python. Target transcription factor-specific and non-specific signals were then quantified as explained in the results section.

### 8 Yeast strain generation and characterization

#### 8.1 Media and growth conditions

Cells were incubated for yeast transformation at 30°C shaking at 250 rpm in yeast extract peptone dextrose (YPD) medium (Sigma-Aldrich) (10 g/L yeast extract, 20 g/L peptone and 20 g/L glucose). Plates contains 20 g/L of agar (10752-36, Alfa).

Promoter library yeast strains were cultured in synthetic complete (SC) medium lacking uracil (1.92 g of yeast synthetic dropout medium supplement without uracil, Y1501-20G, Sigma-Aldrich), 6.7 g Yeast Nitrogen Base (YNB) without amino acids (Y0626-1KG, Sigma-Aldrich), were dissolved in 960 mL of deionized water and autoclaved. After sterilization we added 40 mL 50% filter sterilized glucose solution to a final volume of 1 L. For plate reader experiments of PHO5 promoter library strains we use a different recipe of SC, SC phosphate-free (PF) and SC phosphate-rich (PR) medium, a modified recipe from that described in (Lam et al. (2008)). In this case we add per 1 L, 20 g of glucose, 5.6 g YNB with ammonium sulfate, without phosphates, without sodium chloride (MP 4027-812), 0.79 g Complete Supplement Mixture (CSM) (MPB-114500022) and 0.1 g sodium chloride. The amount of adenine and tryptophan was supplemented over the CSM to final concentrations of 0.13 g/L adenine and 0.1 g/L tryptophan to suppress autofluorescence. SC PF medium contains 0.55 g/L of potassium chloride instead of 1 g/L of monobasic potassium phosphate that contains SC PR medium.

#### 8.2 SC PF medium recipe (per Litre)

| **Ingredient** | **Amount** |
|---|---|
| **Nitogen source** | |
| Ammonium sulfate | 5 mg |
| **Carbon Source** | |
| Dextrose | 20 g |
| **Amino acids** | |
| Adenine | 130 mg |
| L-Arginine | 50 mg |
| L-Aspartic Acid | 80 mg |
| L-Histidine HCl | 20 mg |
| L-Isoleucine | 50 mg |
| L-Leucine | 100 mg |
| L-Lysine HCl | 50mg |
| L-Methionine | 20 mg |
| L-Phenylalanine | 50 mg |
| L-Threonine | 100 mg |
| L-Tryptophan | 100mg |
| Uracil | 20 mg |
| L-Tyrosine | 50 mg |
| L-Valine | 140 mg |
| **Vitamins** | |
| Biotin | 2 µg |
| Calcium Pantothenate | 0.4 mg |
| Folic Acid | 2 µg |
| Inositol | 2 mg |
| Niacin | 0.4 mg |
| P-Aminobenzoic Acid | 0.2 mg |
| Pyridoxine Hydrochloride | 0.4 mg |
| Riboflavin | 0.2 mg |

| | |
|---|---|
| Thiamine Hydrochloride | 0.4 mg |
| **Trace elements** | |
| Boric Acid | 0.5 mg |
| Copper Sulfate | 0.04 mg |
| Potassium Iodide | 0.1 mg |
| Ferric Chloride | 0.2 mg |
| Manganese Sulfate | 0.4 mg |
| Sodium Molybdate | 0.2 mg |
| Zinc Sulfate | 0.4 mg |
| **Salts** | |
| Magnesium Sulfate | 0.5 mg |
| Calcium Chloride | 0.1 mg |
| Potassium chloride | 550 mg |
| Sodium chloride | 100 mg |

Table S7: SC PF medium recipe (per Litre)

#### 8.3 Promoter library and yeast strain construction

Promoter variants were ordered as gene fragments from TWIST Bioscience. The PHO5 promoter library integration cassette consisted of two LYS2 homology arms (each 500 bp), PHO5 promoters variants, yeast codon optimized enhanced green fluorescent protein (yoEGFP), TDH1 terminator sequence (177 bp), and URA3 auxotrophic marker. The minimal CYC1 promoter library integration cassette consisted of two LYS2 homology arms (each 500 bp), CYC1 minimal promoter variants, yoEGFP ENO2 terminator and URA3 auxotrophic marker. All integration cassettes were constructed hierarchically by using modular part plasmids following the YeastToolkit (YTK) workflow (Lee et al. (2015)).

Briefly, multiple YTK part plasmids were assembled into cassette plasmids via BsaI goldengate assembly and then multiple cassette plasmids were assembled to the multi-cassette plasmids via

BsmBI goldengate assembly. Goldengate reactions are prepared as following: add the volume necessary to have 100 ng of each plasmid, except for the backbone that was added 25 ng, 2 µL of 10x T4 ligase buffer (NEB), 1 µL of T4 ligase (NEB), 1 µL of BsaI or BsmBI (NEB), and adding water to final volume of 20 µL. Thermocycler setup: (BsmBI assembly: 45°C for 2 min/BsaI assembly: 42 °C for 2 min, 16°C for 5 min) x 25 cycles, followed by a final digestion step at 60°C for 10 min and a heat inactivation at 80°C for 20 min. To assemble PHO5 promoter library cassette plasmids (from pSC189 to pSC201), assembly connector ConLS part plasmid( pYTK002), PHO5 promoters variant part plasmids (from pSC170 to pSC182), yoEGFP part plasmid (pMC068), TDH1 terminator part plasmid (pYTK056), assembly connector ConRE part plasmid( pYTK072) were assembled into the backbone plasmid (pYTK095) via a BsaI goldengate assembly. PHO5 promoter library integration cassette plasmids (from pML001 to pML013) were constructed by BsmBI assembly of cassettes plasmids (from pSC189 to pSC201) into the backbone plasmid that contains two LYS2 homology arms and URA3 auxotrophic marker (pSC109).

To assemble the minimal CYC1 promoter library cassette plasmids (from pAS009 to pAS016 and from pML022 to pML033), assembly connector ConLS part plasmid( pYTK002), CYC1 minimal promoters variant part plasmids (from pAS001 to pAS008 and from pAS058 to pAS069), yoEGFP part plasmid (pMC068), ENO2 terminator part plasmid (pYTK055), assembly connector ConRE part plasmid (pYTK072) were assembled into the backbone plasmid (pYTK095) via a BsaI goldengate assembly. CYC1 minimal promoter library integration cassette plasmids (from pAS017 to pAS024 and from pML034 to pML045) were constructed by BsmBI assembly of cassettes plasmids (from pAS009 to pAS016 and from pML022 to pML033), into the backbone plasmid that contains two LYS2 homology arms and URA3 auxotrophic marker (pSC109).

For all plasmid cloning we used NEB 10-beta Competent *E. coli* (High Efficiency) (New England Biolabs, Cat C3019H). We followed the transformation protocol from the manufacturer. We conducted bacterial selection and growth in Lysogeny Broth (LB) plates or LB medium at 37°C with supplement of appropriate antibiotics (chloramphenicol 34 µg/mL, ampicillin 100 µg/mL or kanamycin 50 µg/mL). All plasmids were sequence-verified by sanger sequencing (Microsynth Ecoli NightSeq).

The PHO5 promoter library yeast strains were made by transforming *S. cerevisiae* strains BY4741 and ΔPho4 (Yeast Knockout Library, Horizon Discovery Ltd.), with the integration cassettes and we obtained 26 variant strains (from sML034 to sML059). The minCYC1 promoter library integration cassettes were used to transform the parental strains BY4741 and sSC051 (S.C. et al., unpublished data) to obtain 40 variant strains (from sAS001 to sAS008 and from sML073 to sML104). In brief, sSC051 has integrated into HO locus the yeast codon optimized mScarlet-i (yomScarlet-i) under the control of a modified GAL1 promoter with six Zif268 biding sites as described (McIsaac et al. (2012)). Upstream, in the same HO locus, this strain has integrated the artificial transcription factor Z3EV (a fusion of the mouse transcriptional factor Zif268 DNA binding domain, the ligand binding domain of the human estrogen receptor and viral protein 16) that is selectively relocated from the cytoplasm to the nucleus in the presence of $\beta$-estradiol. All promoter library yeast strains were made by homologous recombination of the integration cassettes, into LYS2 locus, using the lithium acetate/polyethylene glycol (PEG) method (Gietz and Schiestl (2007)). For yeast integration all integration cassette plasmids were digested with NotI and after that the DNA was purified (ZYMO DNA clean and concentrator-25 kit). The transformants were selected in SC medium lacking uracil plates at 30°C. We screened transformants for correct integration by colony PCR followed by sanger sequencing of the colony PCR product (Microsynth).

#### 8.4 Yeast characterization

Promoter library strains were grown overnight in 3 ml in SC medium lacking uracil at 30°C shaking at 250 rpm. Cultures were diluted to an optical density (O.D) of 0.175 in fresh SC medium lacking uracil. Cells grown until log phase, OD $\sim$ 0.8, then PHO5 promoter library strains were washed twice in SC phosphate-free (PF) medium and diluted to starting OD of 0.1 – 0.2 in SC PF medium and SC phosphate-rich (PR) medium. Minimal CYC1 promoter strains were diluted to starting OD of 0.1 – 0.2 in SC medium lacking uracil and SC medium lacking uracil with $\beta$-estradiol at 200 nM. Cells were cultured in 96-well plates with clear flat bottom (Nunc) and covered with a gas-permeable membrane (Breathe-Easy membrane, Sigma-Aldrich). OD, GFP and mScarlet was measured every 10 min for 20 – 24 hours on a plate reader (BioTek SynergyMx). 96-well plates were incubated at 30°C with continuous agitation while the plate reading is not measuring. Background of the media was subtracted , then data was normalized dividing by the OD at each time point.

The same growth and inducing conditions were used to induce promoter library for characterization by fluorescence microscopy. The only difference being that 96-well plates with clear conical bottoms (Nunc) were used for growing the yeast for microscope analysis. Strains were imaged at log phase, at an OD of around 0.8, using a Nikon Eclipse Ti fluorescent microscope. Sample visualization was conducted under a 60x objective through the NIS-Elements software (Nikon Instruments). Cells were imaged in bright-field and fluorescence modes. Images were processed and data analyzed using Image J.
